# Inflammatory stimulation redirects SMC1A from speckle-proximal to peripheral chromatin within a functionally compartmentalized genome

**DOI:** 10.64898/2026.07.06.736754

**Authors:** Sofia Papanikolaou, Marianthi Marouli, Anastasia Katifori, Athanasia Mizi, Ariana Cimmarrusti, Despoina Kosmara, Despoina Tsapara, Christos Mantis, Athanasia Stavropoulou, Argyris Papantonis, George Bertsias, Christoforos Nikolaou

## Abstract

Nuclear architectural proteins have multiple regulatory roles, but how they respond to inflammatory stimulation within the three-dimensional nucleus is unclear. Here we show that SMC1A, a core subunit of the cohesin complex, undergoes a striking redistribution upon inflammatory stimulation in human monocytes, between gene populations with distinct sequence composition and nuclear organization. Combining ChIP-seq, RNA-seq and a Chrom3D model of the monocyte nucleus with published chromatin contact, speckle-proximity and mRNA half-life data, we find that this redistribution is associated with multiple levels of gene expression. SMC1A binding is preferentially lost from short, GC-rich genes associated with the nuclear interior and nuclear speckles, whereas binding gains occur at longer, relatively GC-poor genes in more peripheral regions. SMC1A gains are enriched at pre-existing loop anchors coinciding with TAD boundaries, and genes located within loops containing gained anchors are more frequently induced upon inflammation. In parallel, inflammatory stimulation is associated with reduced retention of short, GC-rich introns linked to nuclear speckles, a response that persists upon SMC1A depletion.

These findings suggest that SMC1A redistribution follows a pre-existing genomic organization in which DNA composition, radial positioning and gene architecture are associated with distinct transcriptional and RNA-processing responses to inflammation.

## Introduction

The eukaryotic genome is highly compartmentalized, with chromatin folding hierarchically into loops, topologically associating domains (TADs)^1^, and A/B compartments^2^, while the chromosomes themselves are non-randomly arranged within the three-dimensional space of the nucleus^3,4^. This overarching genome architecture is radially delineated^5,6^. The nuclear periphery lies in proximity to the nuclear lamina^7^ and its genes are enriched in cell-type specific functions that are typically silenced or under tight epigenetic regulation^8^. In contrast, the nuclear interior harbors euchromatic regions, transcription factories, and nuclear speckles (NS)^9^, which represent subnuclear structures linked to constitutively active transcription and RNA processing^10^. Radial genome positioning is known to be correlated with intrinsic gene features including GC content^11^, gene density^12–14^, exon-intron architecture, and splicing complexity^15,16^. For example, GC-rich, and constitutively transcribed genes tend to reside closer to the nuclear center, whereas long, AT-rich are preferentially localized toward the periphery^17^. Across this chromatin milieu, the organization of the eukaryotic genome is remarkably stable, with major rearrangements being confined at the stage of development^18^, while many inducible genes are targeted to the periphery upon activation^19^. A key question is the degree to which this compartmentalized structure may act as a substrate for gene expression programs, and which nuclear components are involved.

Nuclear architectural proteins have multiple roles that extend beyond the maintenance of chromosomal integrity^20^. The high-mobility group protein HMGB2 localizes to nuclear speckles upon replicative senescence^21^, while its relative HMGA1 clusters in dense loci with properties resembling those of constitutive heterochromatin in an oncogene-induced senescence model^22^. The multifunctional insulator protein CTCF creates foci with strong association to splicing^23^ and members of the cohesin complex are actively involved in enhancer-promoter rewiring in RAS-induced senescent fibroblasts^24^. More recently, the cooperative function of CTCF and cohesin was shown to lay down the ground state for chromatin around nuclear speckles^25^.

While depletion of core architectural proteins brings about modest gene expression changes in steady state conditions^26,27^, their role appears to be crucial under stress or development^28–31^. Architectural proteins can fine-tune a pre-existing network of chromatin interactions during cell differentiation events, as we have shown in the case of the nuclear organizer SATB1 during thymocyte maturation^32^.

We have recently shown that SMC1A, a member of the cohesin complex, relocalizes to the enhancers of numerous inflammatory genes upon stimulation of human monocytes^33^. Motivated by this observation, we herein investigate whether this position-dependent gene regulation pattern reflects broader relocalization dynamics within the nucleus.

Here, we show that genes activated under inflammatory conditions are preferentially located at the nuclear periphery and are more likely to gain SMC1A binding. They tend to be long and GC-poor, with proportionally long intergenic distances. SMC1A gains are enriched at pre-existing loop anchors coinciding with TAD boundaries, and genes within loops containing gained anchors are more frequently induced. Conversely, SMC1A is lost from short, GC-rich genes associated with the nuclear interior and speckles, coinciding with reduced retention of short, GC-rich speckle-linked introns, a response that persists after SMC1A depletion. Gene composition and pre-existing genome organization thus provide a structural framework for distinct transcriptional and RNA-processing responses to inflammation.

## Results

### SMC1A associates with distinct gene populations characterized by unique properties prior to and post-inflammatory stimulation

We recently reported a significant relocalization of the nuclear architecture protein SMC1A to the enhancers of immune response genes in human monocytes upon inflammatory, lupus-like stimulation^33^. We asked whether this redistribution extends to different genomic compartments, given that the nuclear periphery is enriched in tissue-specific enhancers^34^. A strong indication for this came from the fact that genes affected by SMC1A knock-down (SMC1Akd) in untreated (442 genes) and stimulated conditions (417 genes) formed two very distinct sets with minimal overlap (Figures 1a, 1b, Supplementary File 1). Both gene sets showed strong functional enrichment for immune-related pathways^33^ but shared only 34 genes (Figure 1b), suggesting that SMC1A is transcriptionally associated with distinct gene sub-populations in untreated and inflamed monocytes.

**Figure 1a.**
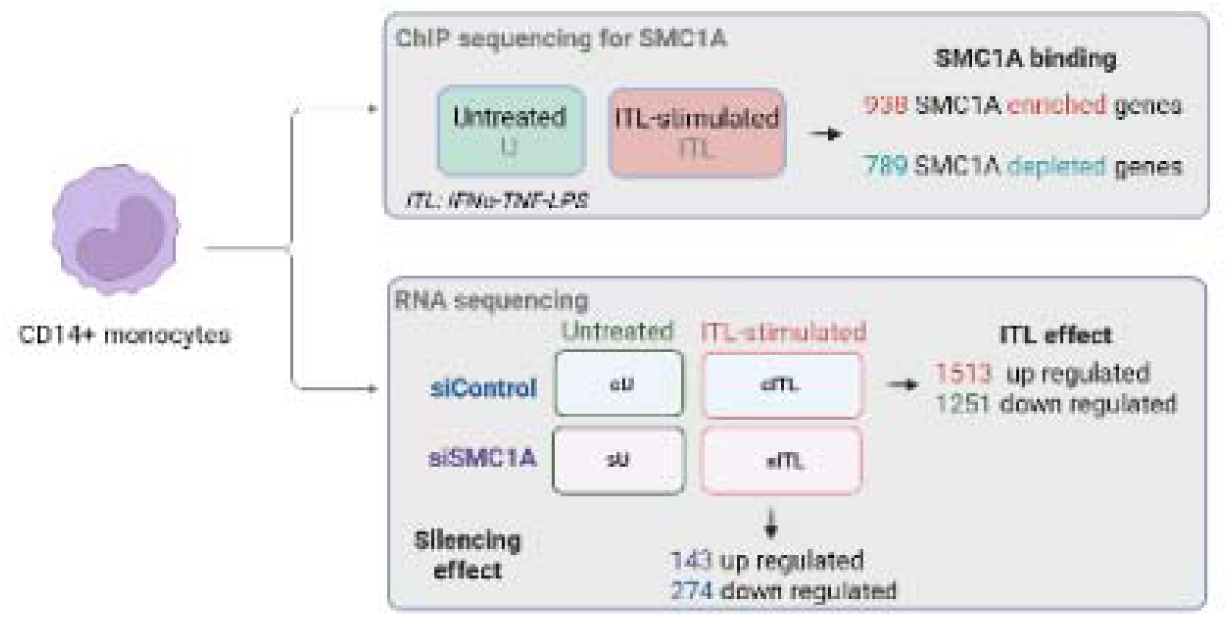
Outline of the experimental conditions analysed in this work. Numbers of affected genes for each comparison appear at the bottom of the panel (Supplementary File 1).

**Figure 1b.**
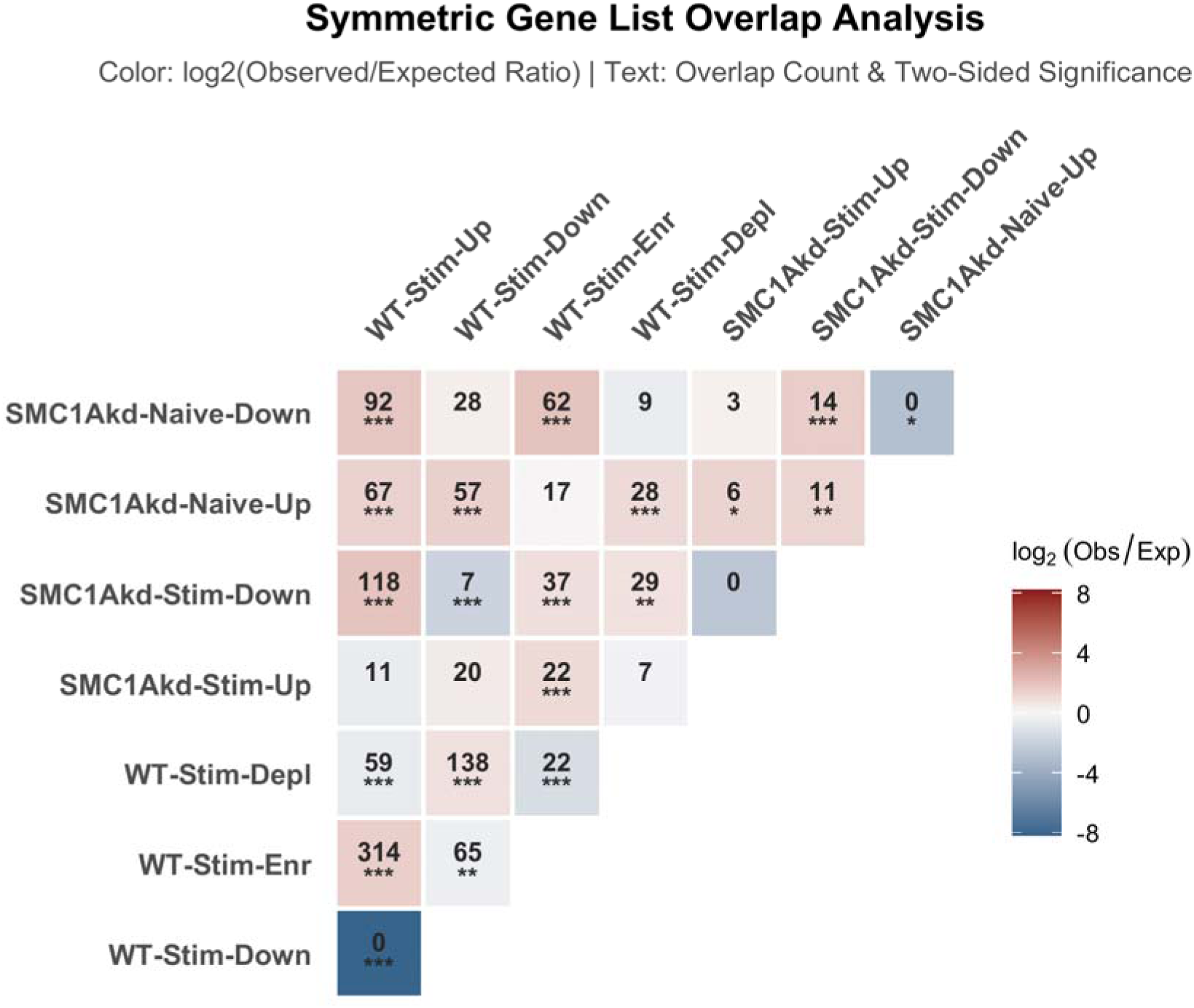
Gene overlaps and relative enrichment (as observed-over-expected ratios) of gene groups described in Figure 1a (stars denote significance of a Fisher’s test at p<0.05 (*), p<0.01 (**) and p<0.001 (***) levels). See also, Supplementary File 1.

When compared against a ChIPSeq experiment against SMC1A in the same (naïve, untreated vs stimulated) conditions, we found differentially activated or repressed genes to be significantly overlapping areas of increased or depleted SMC1A-binding respectively (Figure 1b).

Areas of SMC1A gain/loss also shared some striking differences in terms of DNA composition. Upon inflammatory (ITL) stimulation, SMC1A is strongly depleted from genomic regions with a very high GC% and becomes enriched in regions with lower GC%, close to the genome average (Figure 1c).

**Figure 1c.**
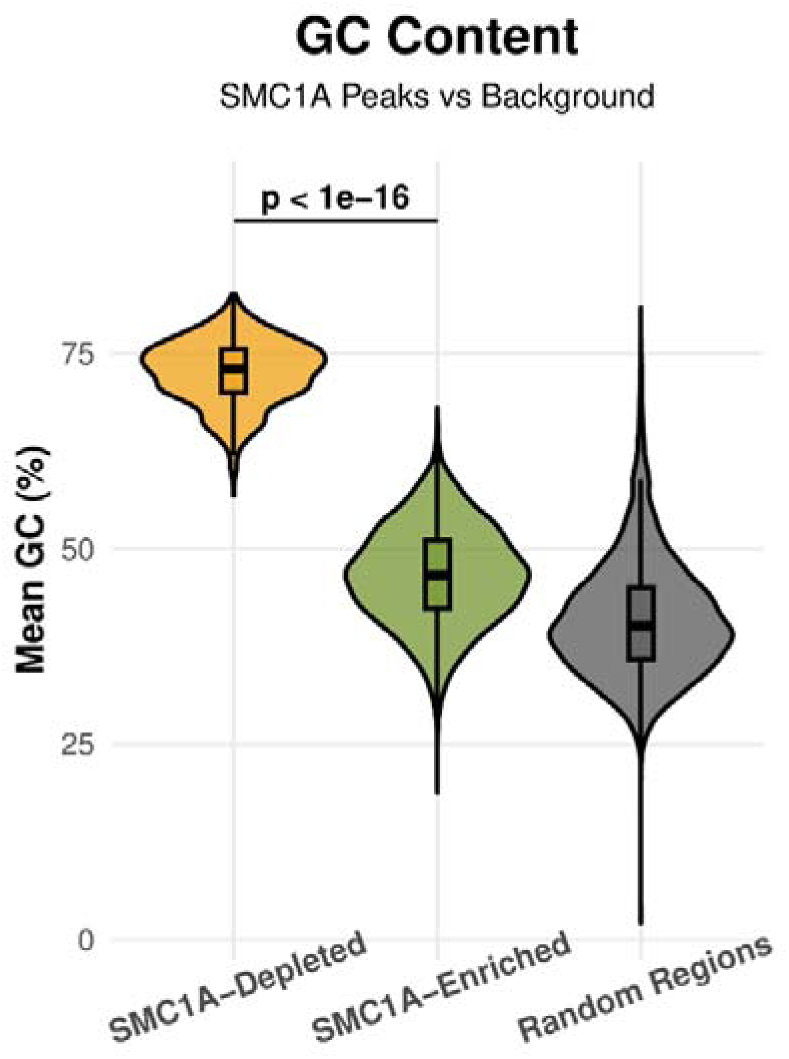
GC% of genomic sites enriched for and depleted of SMC1A upon inflammatory stimulation in human monocytes. Random sites correspond to 5000 randomly selected regions from the human genome, sampling from the size distribution of real sites (p-value of a Wilcoxon Rank-Sum test).

Genomic GC% is associated with numerous structural and functional properties related to chromatin structure, transcriptional activity and DNA replication timing^35^. To identify features related to SMC1A gain/loss beyond GC%, we analyzed a set of structural, evolutionary and sequence properties for all 8 gene groups described in Figure 1a (N=4264, Supplementary File 2, see Methods for details).

Clustering of the 8 gene groups according to their average feature patterns showed a clear separation in two major categories, defined by the average GC% (Figure 1d). Genes depleted of SMC1A and down-regulated in inflammation clustered together at high GC% and short gene lengths with greater than average sequence conservation. On the other hand, genes enriched in SMC1A binding and up-regulated in inflammation clustered around the genome average in terms of GC% and gene length. Notably, genes down-regulated in inflammatory and SMC1Akd conditions showed the most extreme enrichments in alternative splicing events.

**Figure 1d.**
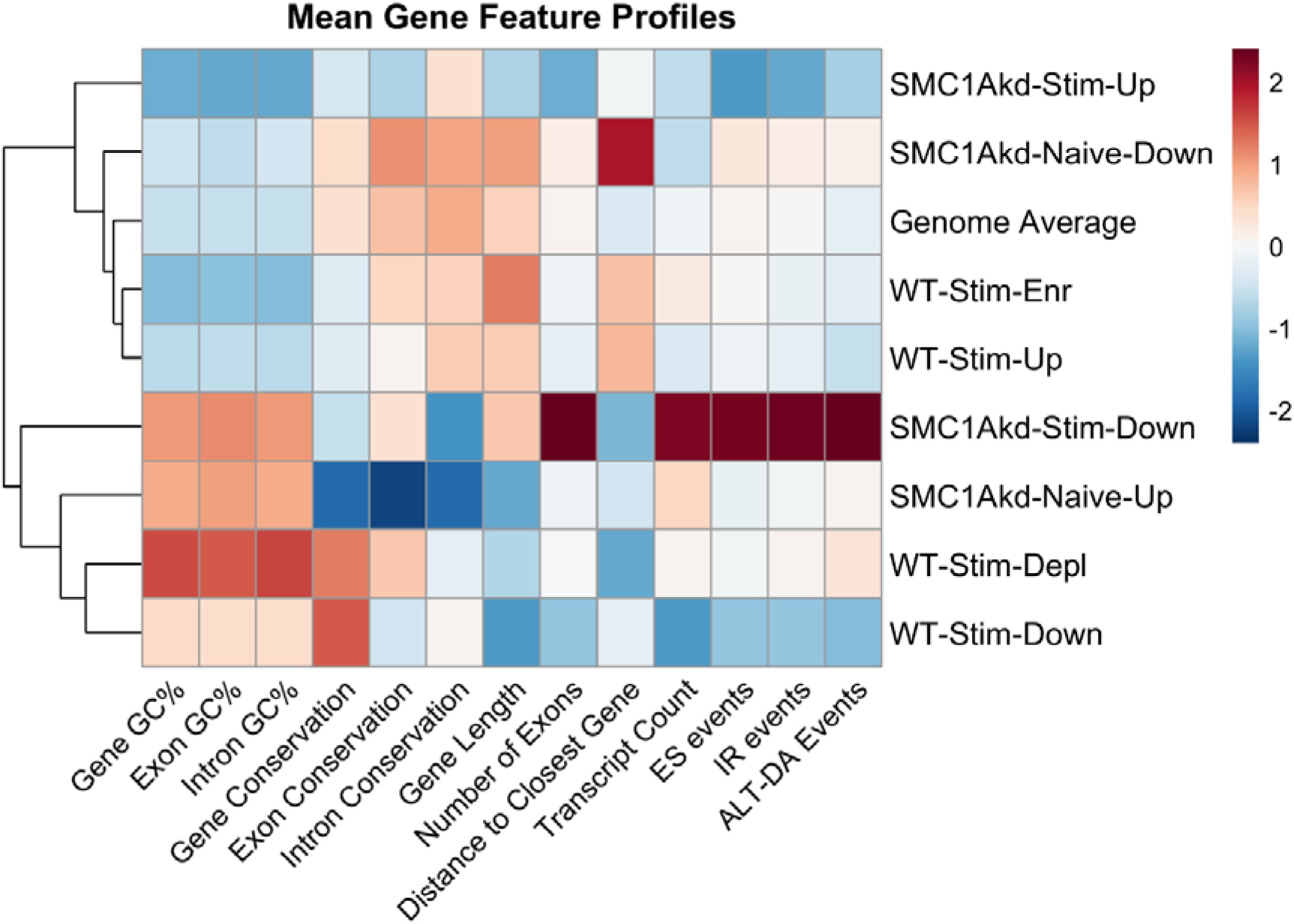
Mean gene profiles for a number of compositional, structural and evolutionary features for the 8 major Gene Classes defined in this study (Supplementary File 2). Every cell in the plot corresponds to a scaled average over all genes in the gene class. (IR: Intron Retention, ES: Exon skipping, ALT-DA: Alternative Donor-Acceptor Events. Clustering is performed using Ward’s D2 to minimize within-cluster variance).

Genes affected by SMC1A knock-down showed equally strong but qualitatively different tendencies. In naïve monocytes, SMC1Akd represses long, GC% poor genes, with average transcriptional complexity (number of transcripts, alternative splicing events) and elevated sequence conservation. These genes also featured the most increased distance to the closest gene compared to the genome average. This is compatible with a preferential association of SMC1A with genes regulated by distant enhancers. This tendency is reversed in stimulated cells, with SMC1Akd severely affecting the expression of densely positioned genes with high GC% and increased transcriptional complexity. Genes that become activated upon SMC1Akd in naïve conditions are also short and GC%-rich but show relatively low transcriptional complexity, with a small number of alternative transcripts and splicing events.

With the aim of disentangling the complexity of these relationships, we used the complete set of gene characteristics to train a Random Forest Model, followed by SHAP (SHapley Additive exPlanations) analysis. GC% and gene length had the highest impact for stimulation-related classes, whereas SMC1Akd-related classes showed lower impact values, with conservation features more predictive in unstimulated and transcriptional and splicing complexity more predictive in stimulated cells (Supplementary Figure 1a; Methods).

An interaction analysis (see Methods) to untangle the contributions of the two perturbations revealed minimal added effect by SMC1Akd, consistent with the SHAP analysis. That is, changes in the combined ITL+SMC1Akd condition, were predominantly attributed to ITL, with only a small number of exceptions (Supplementary Figure 1b).

An analysis of gene profiles, like the one of Figure 1d above, recapitulates the GC%-linked effect of inflammatory stimulation (Supplementary Figure 1c). Inflammation effects are thus predominantly linked to GC%, with SMC1Akd contributing mild additive or, in some cases, synergistic effects associated with transcriptional complexity.

Together, these results indicate that inflammation redistributes SMC1A between two gene populations that differ in sequence composition and gene structure, properties that are known to follow the radial organization of the nucleus.

### SMC1A is depleted from the nuclear center and enriched in the nuclear periphery in inflammatory conditions

Genomic GC% is directly related to the radial positioning of the underlying sequences, with the nuclear center being predominantly occupied by high GC% sequences^16,35^. We thus hypothesized that SMC1A may be selectively evicted from the center and redistributed to more peripheral genome regions upon inflammatory stimulation. To test this, we used a Hi-C map of primary monocytes (ENCODE ENCSR236EYO) and Chrom3D^36^ to create a 3D model of the monocyte nucleus (Figure 2a, see Methods).

**Figure 2a.**
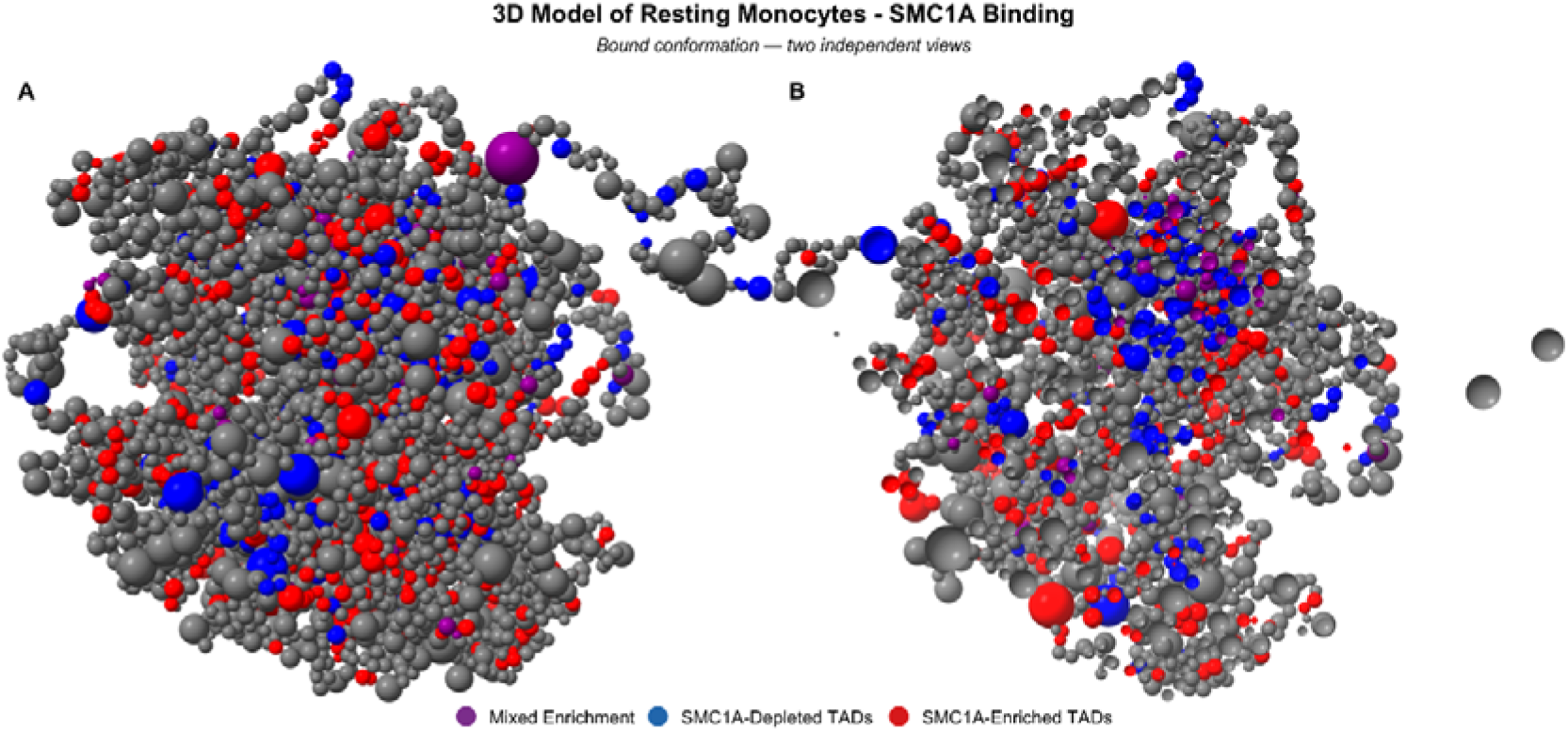
Chrom3D^36^ model representations of naïve monocytes using ChimeraX^37^. Each bead corresponds to a TAD defined from the Hi-C data using HiC-Pro^38^. Red beads correspond to TADs enriched in SMC1A binding and blue beads correspond to TADs with depleted SMC1A binding upon inflammatory stimulation. Purple beads correspond to combined enrichment. Grey beads suggest unaffected TADs.

To better assess the radial position preferences of our gene groups we used a binning approach by dividing the genome 3D-space into 5 radial zones of increasing distance from the center of the nucleus. We then calculated the relative enrichment, as observed-over-expected ratio, of genes associated with SMC1A binding, the inflammatory response and SMC1Akd in each of the 5 zones and assessed its significance based on a Fisher’s test (see Methods). Genes that lost SMC1A binding upon inflammatory stimulation showed the strongest radial preference, being enriched in the two most central zones and depleted from the two most peripheral ones (Figure 2b).

**Figure 2b.**
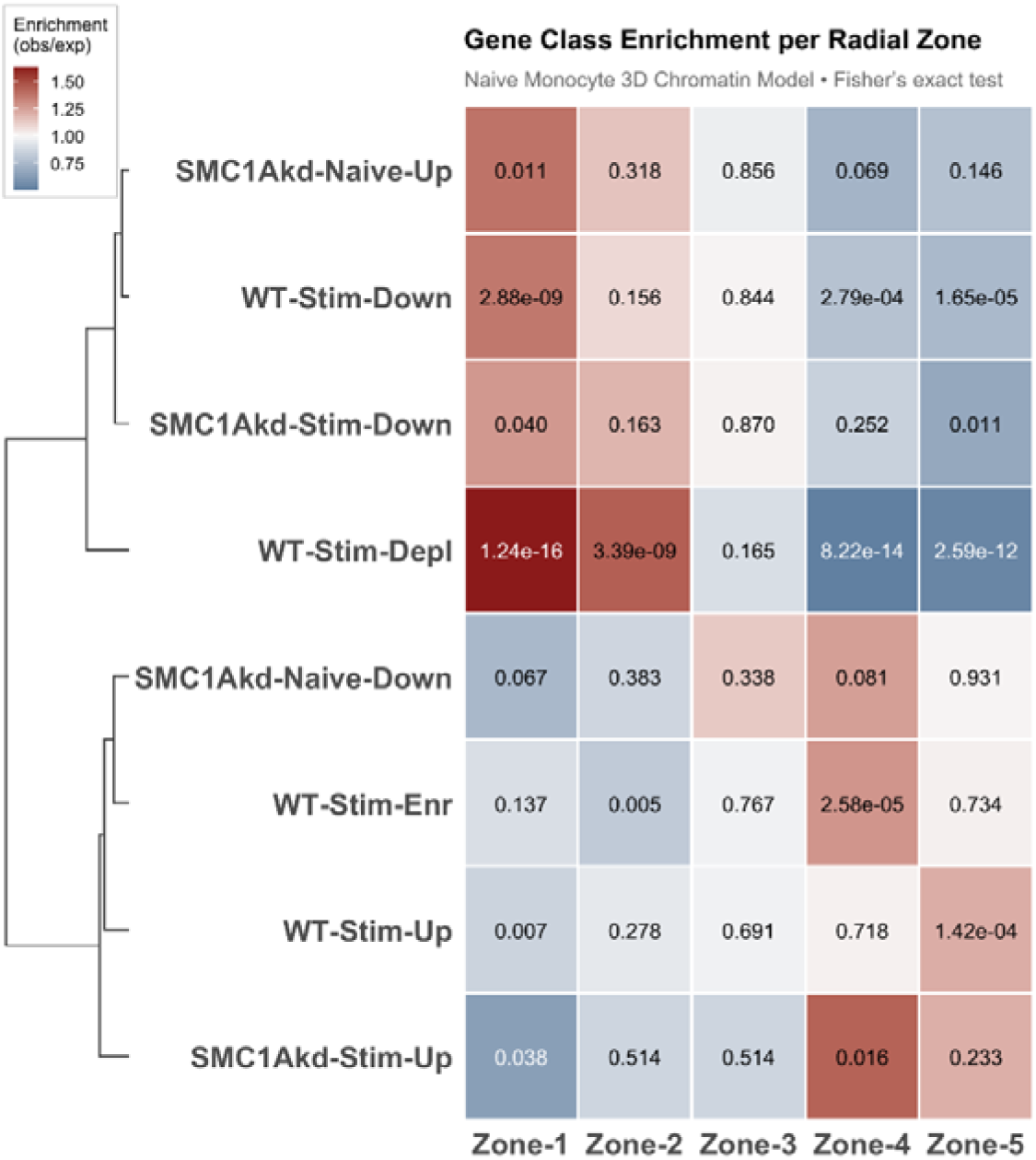
Relative enrichment of gene classes in increasing radial distance zones based on our Chrom3D^36^ model of naïve monocytes. Color coding denotes fold-enrichment. Values inside the cells correspond to p-values of a two-sided Fisher’s test.

Genes down-regulated by inflammation followed the same pattern, whereas genes up-regulated by inflammation showed the opposite trend, being depleted from the central zone and enriched in the most peripheral one. Genes that gained SMC1A binding showed a more moderate shift, with enrichment in zone 4 and depletion from zone 2. Accordingly, genes that lost SMC1A were positioned closer to the nuclear center than random genes, regardless of their expression state. Among genes that gained SMC1A, it was the subset induced by inflammation that was positioned furthest from the center (Supplementary Figure 2).

Clustering of gene groups by radial enrichment mirrored the feature-based clustering (Figure 1d), indicating that radial position covaries with gene structural and evolutionary properties beyond GC%.

Although the model represents the basal monocyte state, genes were differentially modulated by stimulation in a spatially dependent manner. This finding is in agreement with recent works that propose revisiting the view of the nuclear periphery as an overall repressive environment and consider it, instead, a facultatively active transcriptional niche^39,40^. Differential chromatin tethering by architectural proteins such as CTCF and cohesin has been shown to play a direct role in the process of activation by promoting the focal detachment of chromatin from the nuclear lamina^41^.

Because the model represents naïve monocytes, we asked how stable radial positions are across cell types and methods. We first used GPSeq scores obtained with an orthogonal, crosslinking-free protocol^5^. Average GPSeq scores from HAP1 cells (myeloid origin) correlated significantly with our Chrom3D radial distances (Supplementary Figure 3a). Smaller, yet significant correlations were obtained for GPSeq scores from lymphoid GM06990 cells (Pearson’s r=0.268, p-value=0.00012). We also compared zonal assignments with a similarly obtained Chrom3D model for K562 cells, from a recent work^16^. Zonal assignments of shared genes (N=10137) were strongly associated, suggesting consistent radial distributions across cell types (Supplementary Figure 3b, χ^2^ test p-value<10^-16^). Radial positions were thus largely conserved across cell types and methods, although we cannot rule out some gene relocalization upon inflammation. These comparisons suggest that broad radial organization is shared across the examined cell types and approaches. However, as our model represents resting monocytes, we cannot determine whether individual genomic regions change their radial positioning upon stimulation.

We next examined the radial distribution of individual sites. Each differential SMC1A, H3K27ac and ATAC-seq site was assigned to the Chrom3D bead containing it, and its enrichment in each zone was calculated against all tested sites of the same assay (Figure 2c). Sites that lost SMC1A binding upon stimulation were over-represented in the central zones and depleted from the periphery (observed-over-expected ratio 1.83 in zone 1 and 0.44 in zone 5), while sites that gained SMC1A showed the opposite, although weaker, trend (0.81 and 1.14). Changes in H3K27ac followed the same pattern (1.47 to 0.34 for sites with reduced and 0.61 to 1.23 for sites with increased signal). Changes in chromatin accessibility behaved differently: sites that lost accessibility, almost twice as many as those that gained it, were depleted from the center and enriched at the periphery (0.52 and 1.46), while sites with increased accessibility showed only a slight peripheral preference (0.99 and 1.15). Inflammatory stimulation is thus associated with a net shift of SMC1A binding and H3K27ac from the nuclear center towards the periphery, which is not mirrored by changes in chromatin accessibility.

**Figure 2c.**
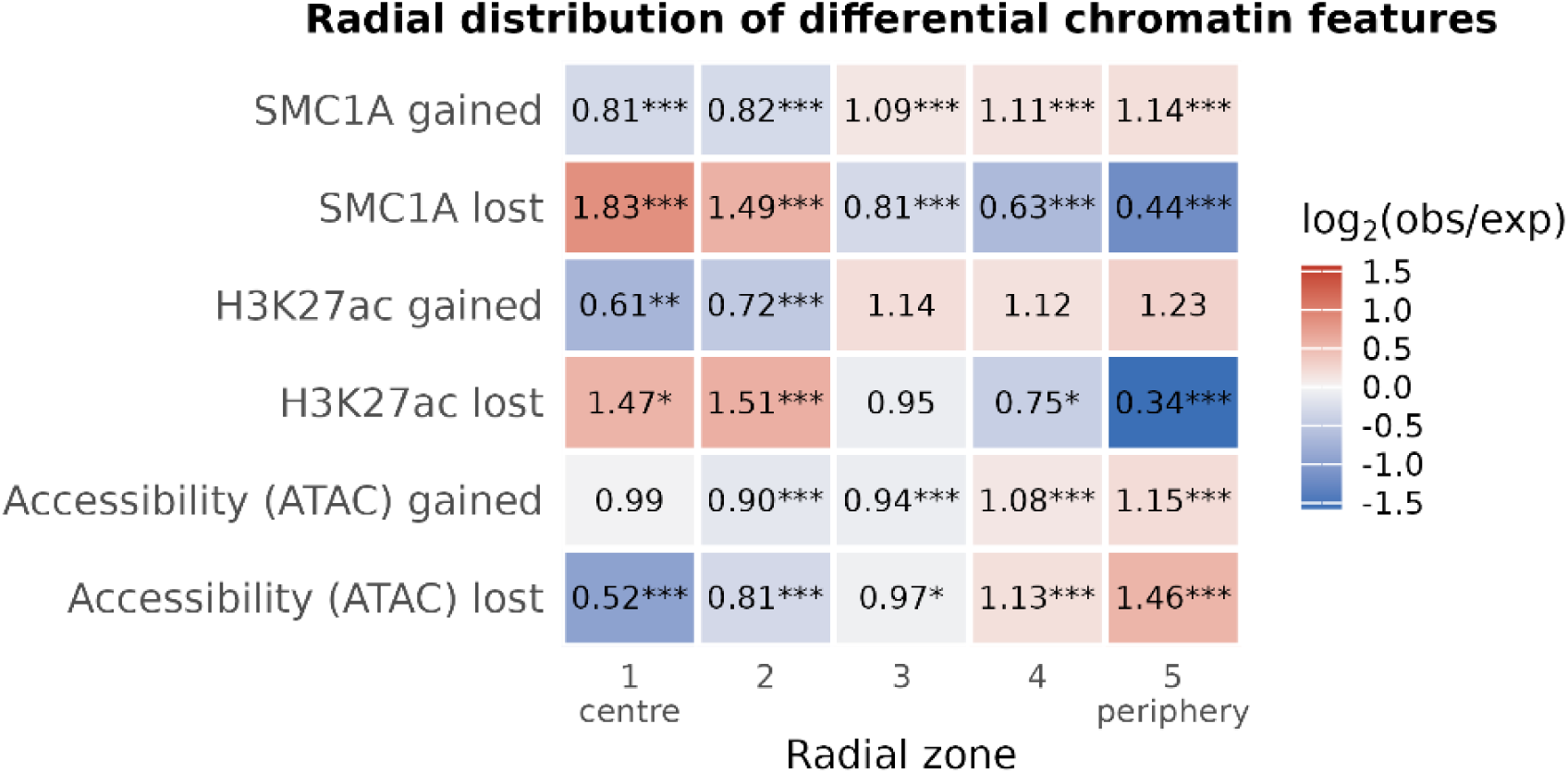
Radial distribution of differential SMC1A, H3K27ac and ATAC-seq sites (stimulated versus untreated monocytes). Each site was assigned to the Chrom3D^36^ bead containing its midpoint; the five zones are equal-width bins of the normalized radial distance. Color scale is logarithmic; values in cells are observed-over-expected ratios.

This radial pattern was closely associated with gene composition. Within each of the five zones, genes that lost SMC1A had a higher GC% than genes that gained it (median 0.57, 0.58, 0.53, 0.53 and 0.44 versus 0.49, 0.46, 0.44, 0.42 and 0.41 from zone 1 to zone 5; Wilcoxon p ≤ 10⁻⁴ in every zone). Genes that gained SMC1A were also longer than the remaining genes in every zone (median 50–116 kb versus 16–35 kb; Figure 2d). Thus, SMC1A binding changes are associated with distinct gene properties even within the same radial zone. As GC-rich genes predominate in central regions and longer genes are more frequent towards the periphery, gene composition likely contributes to the observed radial distribution of SMC1A gains and losses.

**Figure 2d.**
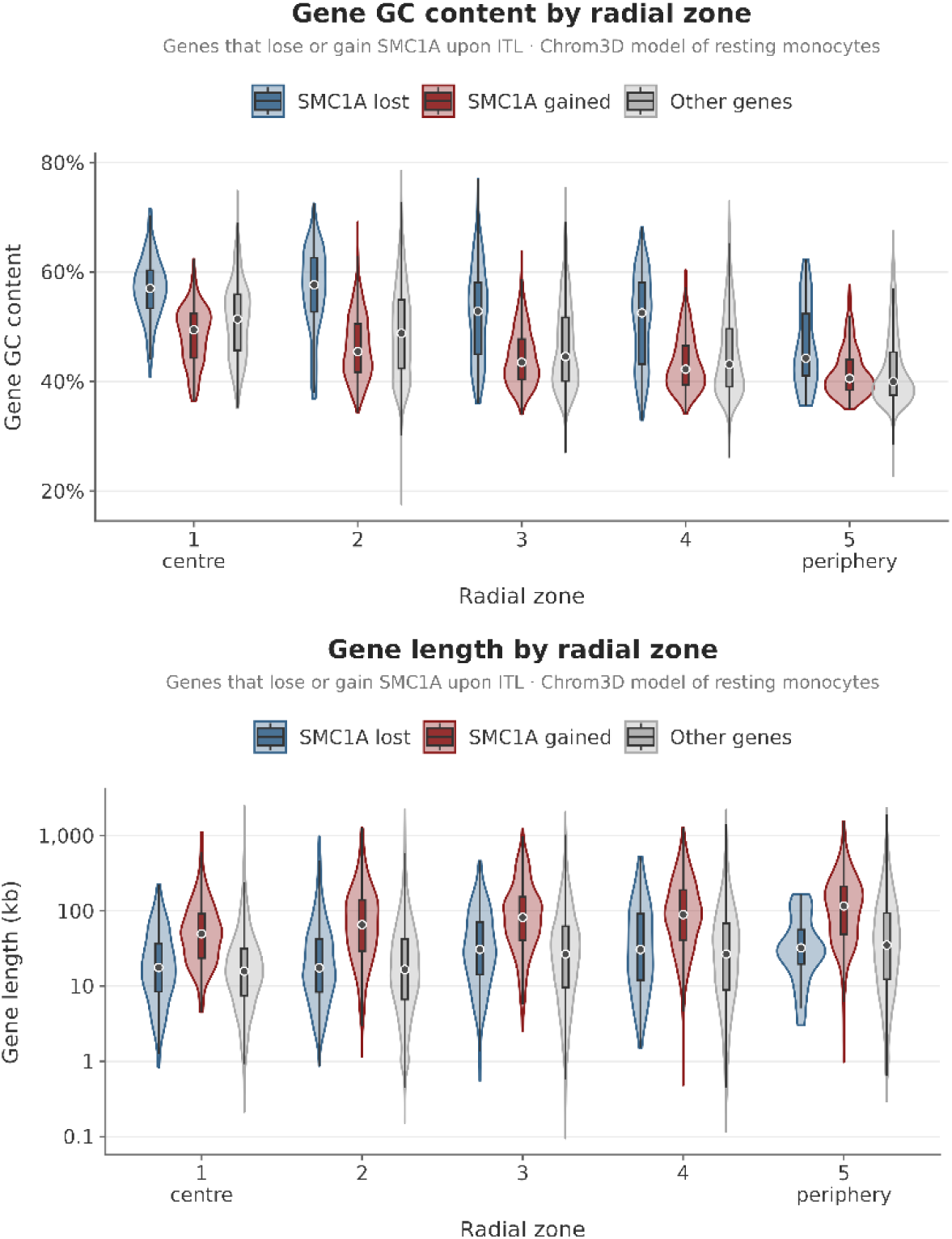
GC% (top) and length (bottom) of genes that lose or gain SMC1A upon inflammatory stimulation, compared to all other genes, in each of the five radial zones. A gene was assigned to a group if the midpoint of a differential SMC1A site (p<0.01, |fold|≥1) lay within the gene ±1 kb. P-values of Wilcoxon Rank Sum tests.

Cohesin is best known for its role in chromatin loop organization. We therefore examined whether inflammation-associated changes in SMC1A occupancy were related to pre-existing chromatin loops and topologically associating domains (TADs), using Hi-C data from resting human monocytes (see Methods). Most SMC1A sites overlapped chromatin loop anchors, irrespective of their response to stimulation. Sites gaining SMC1A were preferentially enriched at loop anchors coinciding with TAD boundaries (19.0% versus 12.0% of unchanged sites; odds ratio [OR] = 1.72, 95% confidence interval [CI] = 1.60–1.86), whereas no enrichment was observed at loop anchors outside TAD boundaries (55.0% versus 55.8%; Figure 2e (left)). In contrast, sites losing SMC1A were under-represented at loop anchors and more frequently located outside both loop anchors and TAD boundaries (46.5% versus 29.0%; OR = 2.13, 95% CI = 1.88–2.41). These findings reveal an asymmetric relationship between SMC1A redistribution and pre-existing chromatin architecture, with binding gains preferentially associated with loop anchors at TAD boundaries and binding losses enriched outside these architectural features.

**Figure 2e.**
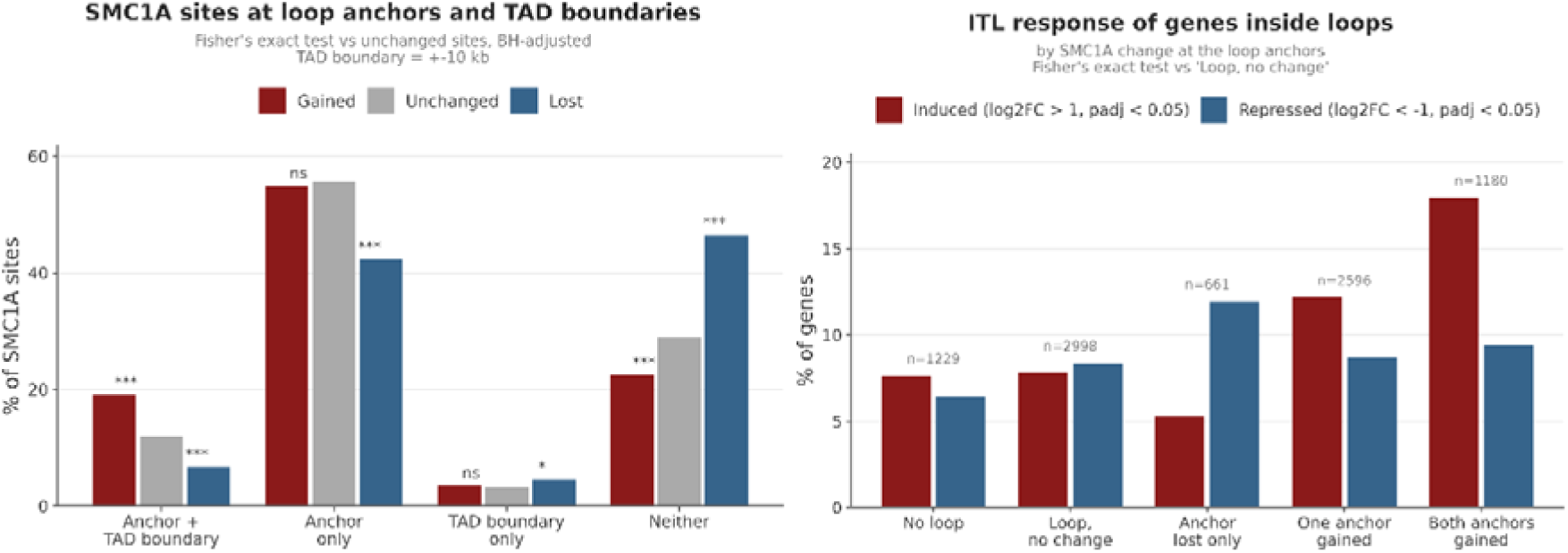
(left) Distribution of SMC1A-binding sites gained, lost or unchanged following inflammatory stimulation across four mutually exclusive genomic categories: loop anchors coinciding with TAD boundaries, loop anchors outside TAD boundaries, TAD boundaries outside loop anchors, and regions overlapping neither feature. Chromatin loops and TADs were identified from Hi-C data of resting human monocytes. Sites were considered anchor-associated when overlapping a loop anchor and boundary-associated when their midpoint was located within 10 kb of a TAD boundary. Bars indicate the percentage of sites within each SMC1A-binding class. Statistical significance was assessed using Fisher’s exact tests comparing gained or lost sites with unchanged sites in each category, with Benjamini-Hochberg correction across eight comparisons (***adjusted P < 0.001; *adjusted P < 0.05; ns, not significant). (right) Percentage of genes induced or repressed following inflammatory stimulation according to the SMC1A-binding status of pre-existing chromatin loop anchors. Genes were assigned to loops when their entire gene body was contained within the genomic interval spanned by the two anchors. Genes overlapping multiple loops were assigned to mutually exclusive categories according to the highest-priority loop status: both anchors gained SMC1A, one anchor gained SMC1A, lost anchor only, or no differential SMC1A binding at loop anchors. Genes outside all loops were classified separately (*No loop*). Gene induction and repression were defined as adjusted P < 0.05 with log2 fold change > 1 or < −1, respectively, in ITL-stimulated versus untreated siControl monocytes. Statistical comparisons were performed using Fisher’s exact tests against genes within loops without differential SMC1A binding at their anchors (*Loop, no change*). Numbers above the bars indicate the number of genes in each category.

We next investigated whether SMC1A redistribution at loop anchors was associated with inflammatory transcriptional responses. Genes located within loops containing SMC1A-gained anchors were more frequently induced following ITL stimulation, with induction increasing according to the number of gained anchors (7.8%, 12.2% and 18.0% for loops with zero, one or two gained anchors, respectively; OR = 1.64 and 2.59; Figure 2e, right). Conversely, genes located within loops containing SMC1A-lost anchors were more frequently repressed (12.0% versus 8.4%; OR = 1.49; Figure 2e, right).

As chromatin loops and TAD boundaries were identified in resting monocytes, these analyses do not establish whether chromatin architecture changes upon stimulation. Rather, they suggest that SMC1A gains at pre-existing loop anchors are associated with increased inflammatory gene induction within the genomic regions spanned by these loops.

### SMC1A binding is lost from speckle-proximal regions upon inflammatory stimulation

The higher GC% and central positioning of genes that lose SMC1A binding upon stimulation prompted us to examine their association with nuclear speckles (NS). CTCF and members of the cohesin complex have been shown to associate with the, centrally-located, NS and to directly influence splicing patterns^23,25^, while CTCF has been shown to dissociate from NS upon stress and during differentiation of neuronal stem cells^42^. To examine whether SMC1A binding changes follow a similar pattern, we obtained TSA-Seq^43^ data for the NS marker SON in a myeloid cell line (K562)^16^, as the closest available model to human monocytes. SMC1A-enriched and -depleted regions were mapped against TSA-Seq percentiles (100% = closest to speckles; Methods).

Genes that lost SMC1A binding were significantly closer to NS than those that gained it, and showed similar proximity to a set of constitutively NS-associated genes^44^ (Figure 3a). By contrast, genes that gained SMC1A showed NS proximity similar to random genes. Notably, genes up-regulated upon SMC1A knockdown in untreated monocytes were not enriched near speckles, despite their relatively central positioning. These genes were generally short and GC-rich, but lacked the transcriptional complexity and alternative splicing characteristics of NS-associated genes (see Figure 1d).

**Figure 3a.**
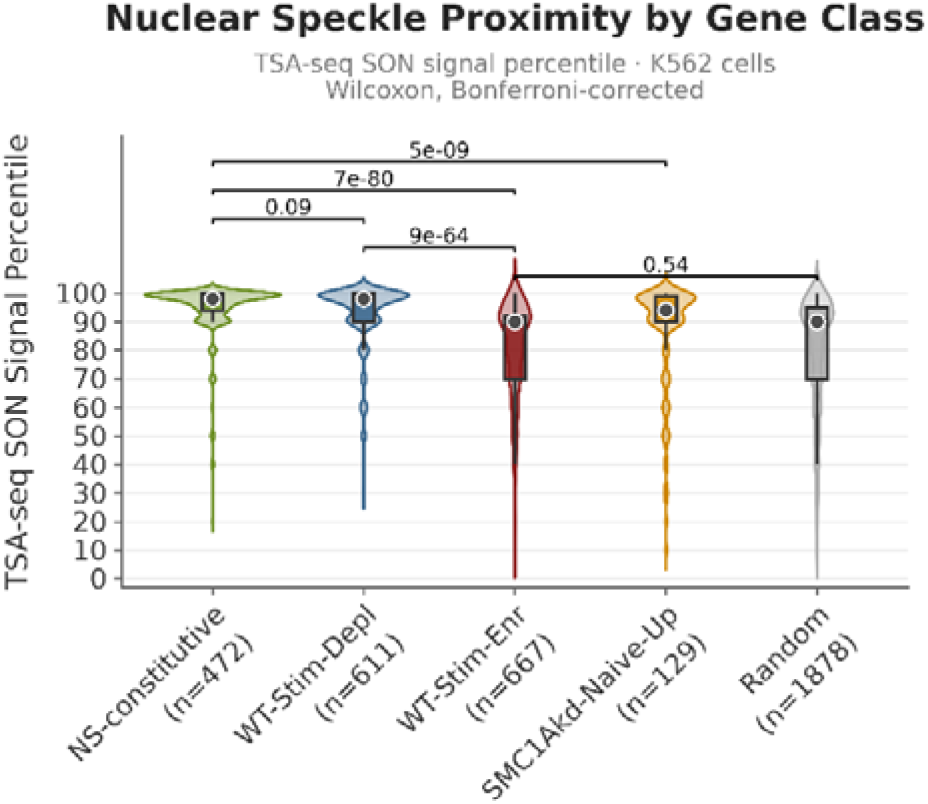
Nuclear Speckle proximity of SMC1A-depleted genes (blue) compared to SMC1A-enriched (red) and a set of NS-constitutive genes^44^. Proximity was quantified as quantile score of a TSA-SON experiment in K562 cells, obtained from^16^, with 100% percentile corresponding to minimal distance from a nuclear speckle. P-values from pairwise Wilcoxon Rank Sum tests.

We next compared our gene groups with gene sets constitutively-, transiently- or non-associated with NS, defined by in situ reverse transcription^44^. Genes losing SMC1A binding upon inflammation were among the most enriched in NS-associated transcripts (Figure 3b). Genes repressed by SMC1Akd showed opposite patterns in the two conditions: they were the most depleted from NS in unstimulated monocytes, but the most NS-enriched in activated monocytes. In agreement with the independent SON data, SMC1Akd up-regulated genes in untreated cells showed no enrichment among NS-associated transcripts.

**Figure 3b.**
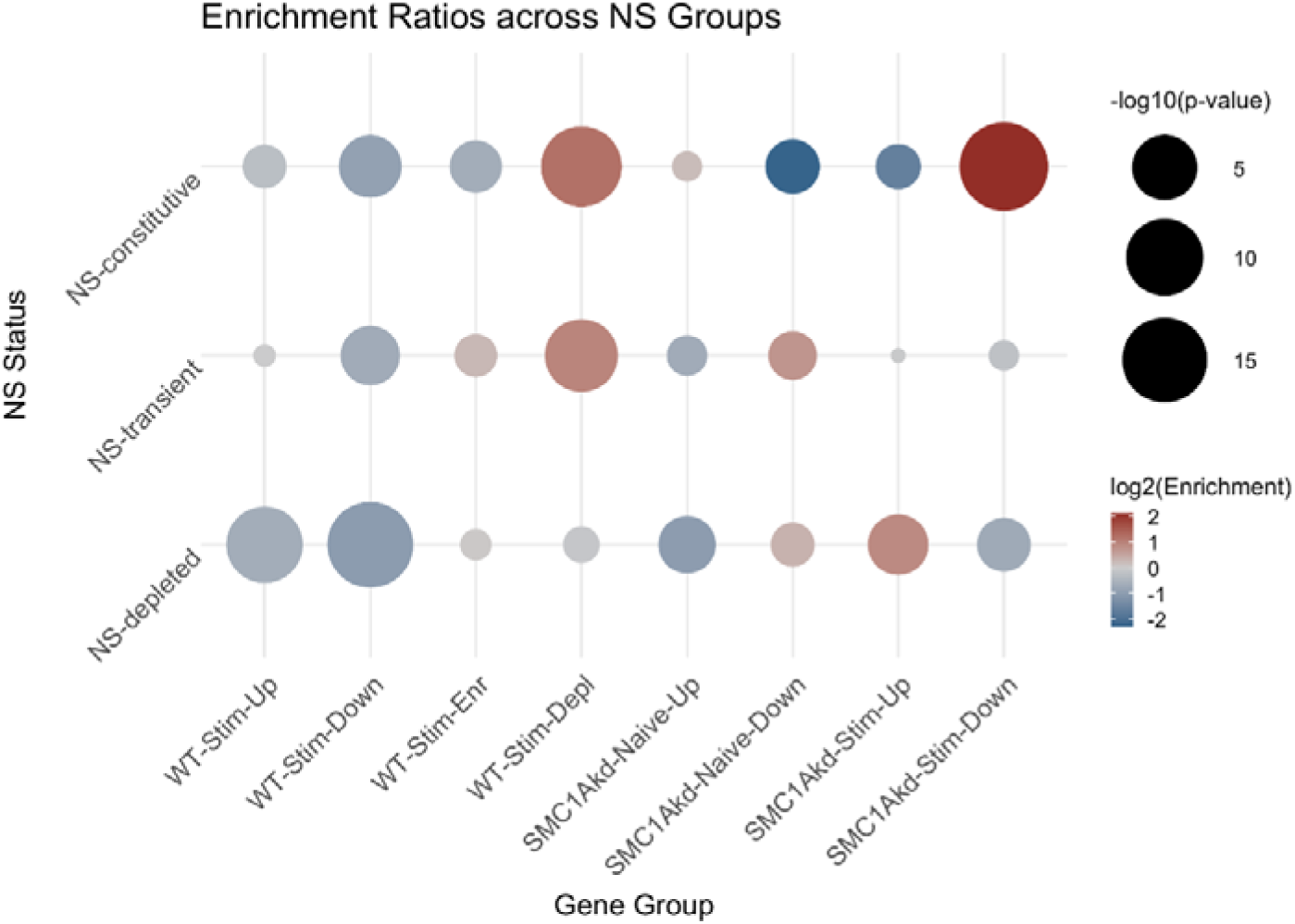
Relative enrichment of inflammatory or SMC1Akd gene groups among Nuclear Speckle associated genes as defined through an in situ reverse transcription approach^44^. Enrichments were calculated as observed over expected ratios of gene overlaps. P-values were assessed on the basis of Fisher’s exact tests (BH-adjusted for multiple comparisons).

Together, these observations suggest that inflammatory stimulation preferentially reduces SMC1A binding at genes associated with nuclear speckles, whereas genes affected by SMC1A depletion show different speckle-association patterns depending on the inflammatory state.

Genes were also compared with published SPIN states in K562 cells^45^ (see Methods). Genes that lose SMC1A binding and/or are repressed upon stimulation were enriched in speckle-associated states, whereas genes with elevated expression and SMC1A binding were enriched at increasing distances from the nuclear center but not in lamina-associated states (Supplementary Figure 3c).

### SMC1A dissociation from nuclear speckles coincides with a reduction in intron retention events

We next compared alternative splicing in stimulated versus unstimulated monocytes, both qualitatively (percentages of alternative splice events) and quantitatively (changes in Percent Spliced In, PSI; see Methods).

Upon stimulation, intron retention (IR) events were predominantly reduced (399 introns with decreased versus 261 with increased retention, |ΔPSI| ≥ 0.15), whereas exon skipping (ES) events changed in the opposite direction (283 versus 359; OR = 1.94, Bonferroni-adjusted p = 1.5 × 10⁻⁸; Figure 3c).

**Figure 3c.**
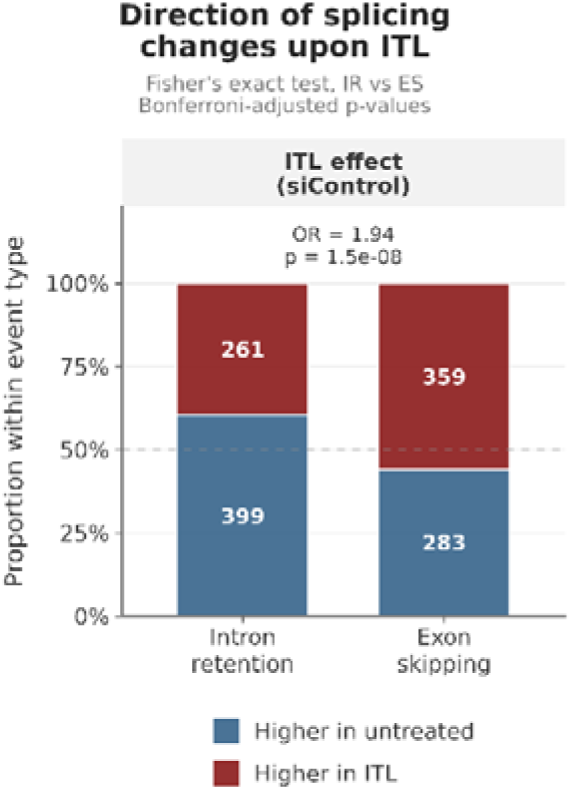
Direction of intron retention (IR) and exon skipping (ES) changes upon ITL stimulation. Bars show the proportion of events with higher inclusion in untreated (blue) or ITL-stimulated (red) monocytes among events with |ΔPSI|≥0.15; numbers of events are shown inside the bars. Odds ratio and p-value of a Fisher’s exact test comparing the direction of IR and ES events (Bonferroni-adjusted shown here and in Figure 4a).

Introns that lost retention upon stimulation were shorter (p = 1.2 × 10⁻¹⁴), more GC-rich (p = 7.3 × 10⁻⁸) and had a smaller exon–intron GC% differential (p = 4.7 × 10⁻⁵) compared with introns that gained retention (Figure 3d, see Methods). These characteristics are consistent with intron-definition splicing and are commonly found in speckle-associated genes^16^.

**Figure 3d.**
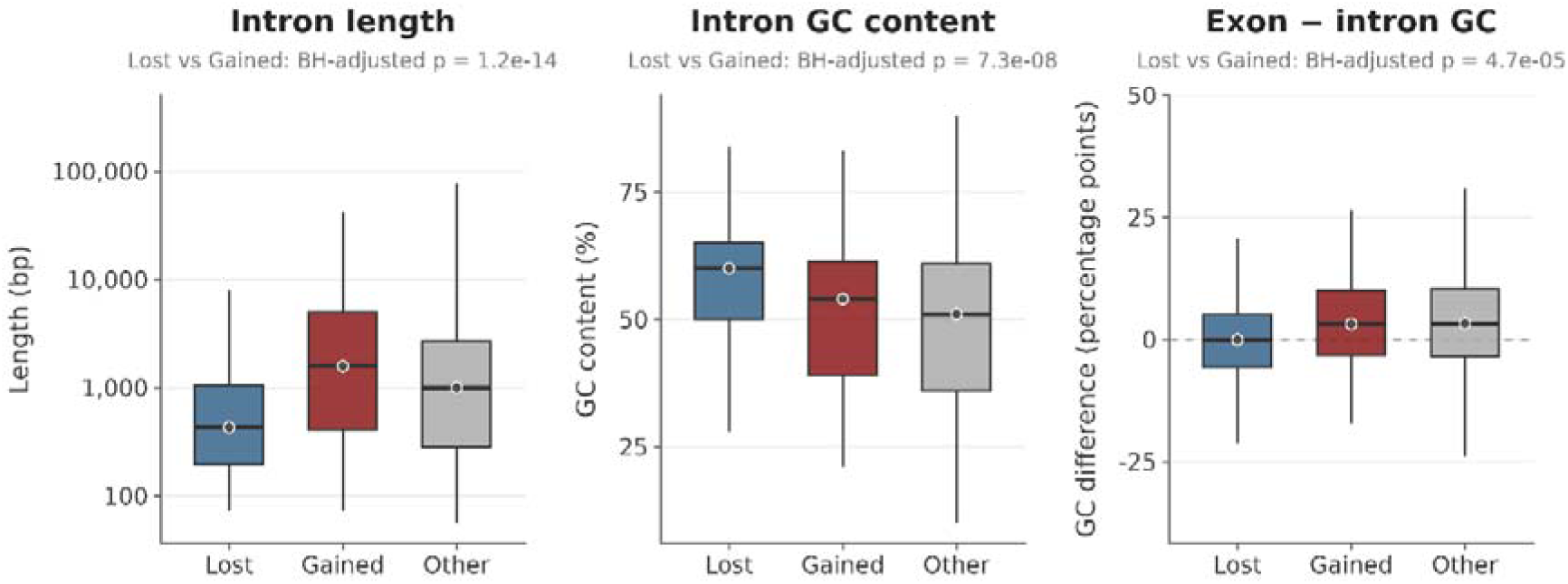
Length (left), GC content (middle) and exon-intron GC difference (right; mean GC% of the two flanking exons minus intron GC%) of introns that lose (Lost) or gain (Gained) retention upon ITL, compared with all other measured introns (Other). Per-intron values obtained from^16^. P-values of Wilcoxon Rank Sum tests, Lost versus Gained, BH-adjusted.

We therefore examined whether introns with reduced retention were associated with nuclear speckles. Their host genes showed higher TSA-SON proximity scores than genes containing introns with retention (p = 7.9 × 10⁻⁵) or genes without IR changes (p = 1.1 × 10⁻¹⁴; Figure 3e).

**Figure 3e.**
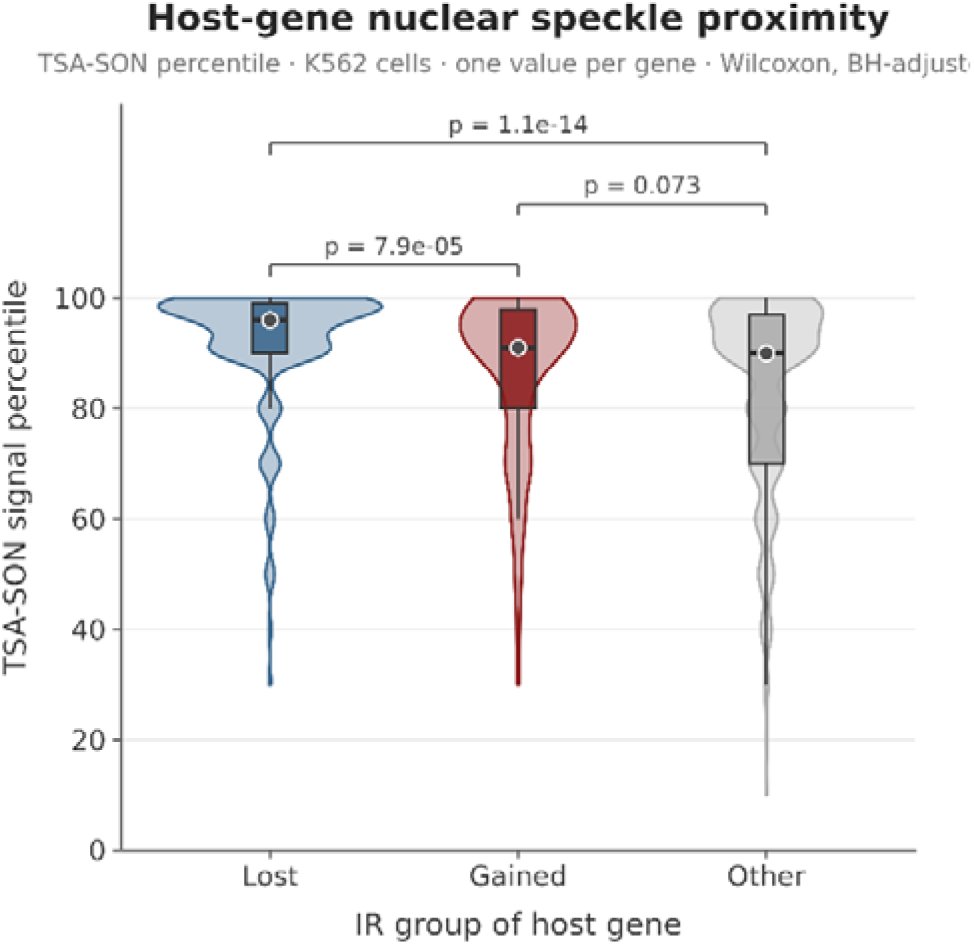
Nuclear speckle proximity of the genes hosting the introns shown in d, measured as TSA-SON percentile in K562 cells^16^; (100th percentile, closest to speckles). Each gene is represented once: genes with at least one intron losing retention upon ITL (Lost), with at least one intron gaining retention (Gained), or with measured introns but no change (Other); genes with introns in both the Lost and Gained groups were excluded. P-values of Wilcoxon Rank Sum tests between each pair of groups, BH-adjusted.

In agreement with this observation, among introns also measured by APEX-seq in HEK293 cells^15^, introns more strongly enriched near SRSF1-marked speckles showed larger losses of retention upon stimulation (ρ = −0.19, p = 4.2 × 10⁻⁵, n = 446), whereas no such relationship was found for lamina-proximal (LMNA) introns (ρ = 0.09, p = 0.12, n = 331; Figure 3f).

**Figure 3f.**
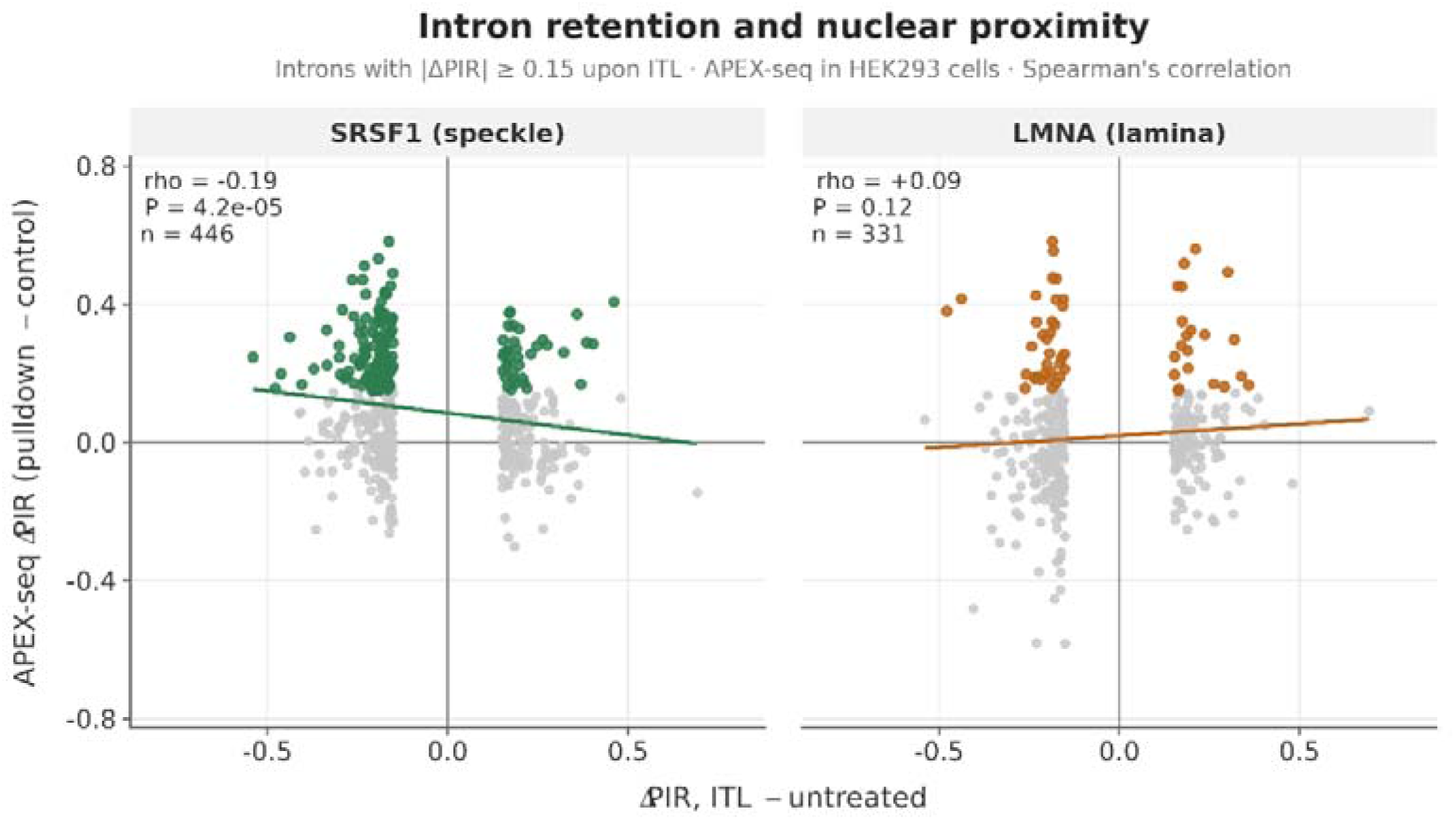
Change in intron retention upon ITL (ΔPIR, ITL − untreated) against the enrichment of the same introns in APEX-seq data of HEK293 cells^15^, for speckle (SRSF1, left) and lamina (LMNA, right) proximity labeling (APEX-seq ΔPIR, pulldown − nuclear control). Introns with |ΔPIR|≥0.15 upon ITL that are measured in the APEX-seq data are shown; coloured points are introns enriched in the corresponding proximity labeling (≥0.15). Lines show linear fits; ρ and P from Spearman’s rank correlation.

Together, these results indicate that inflammatory stimulation preferentially reduces retention of short, GC-rich introns associated with speckle-proximal RNA populations. These introns share compositional characteristics with genes that lose SMC1A binding upon stimulation.

### Inflammation-associated intron retention changes persist upon SMC1A silencing

We next asked whether the reduction in intron retention upon inflammatory stimulation is maintained at reduced SMC1A levels. In SMC1A-silenced monocytes, ITL stimulation still reduced intron retention, although the imbalance between IR and ES events was attenuated (OR = 1.28, p = 0.117, versus 1.94 in control cells; Figure 4a). SMC1A knock-down alone also reduced IR in untreated (OR = 1.68, p = 1.7 × 10⁻⁴), but not in stimulated monocytes (OR = 1.12, p = 1; Figure 4a).

**Figure 4a.**
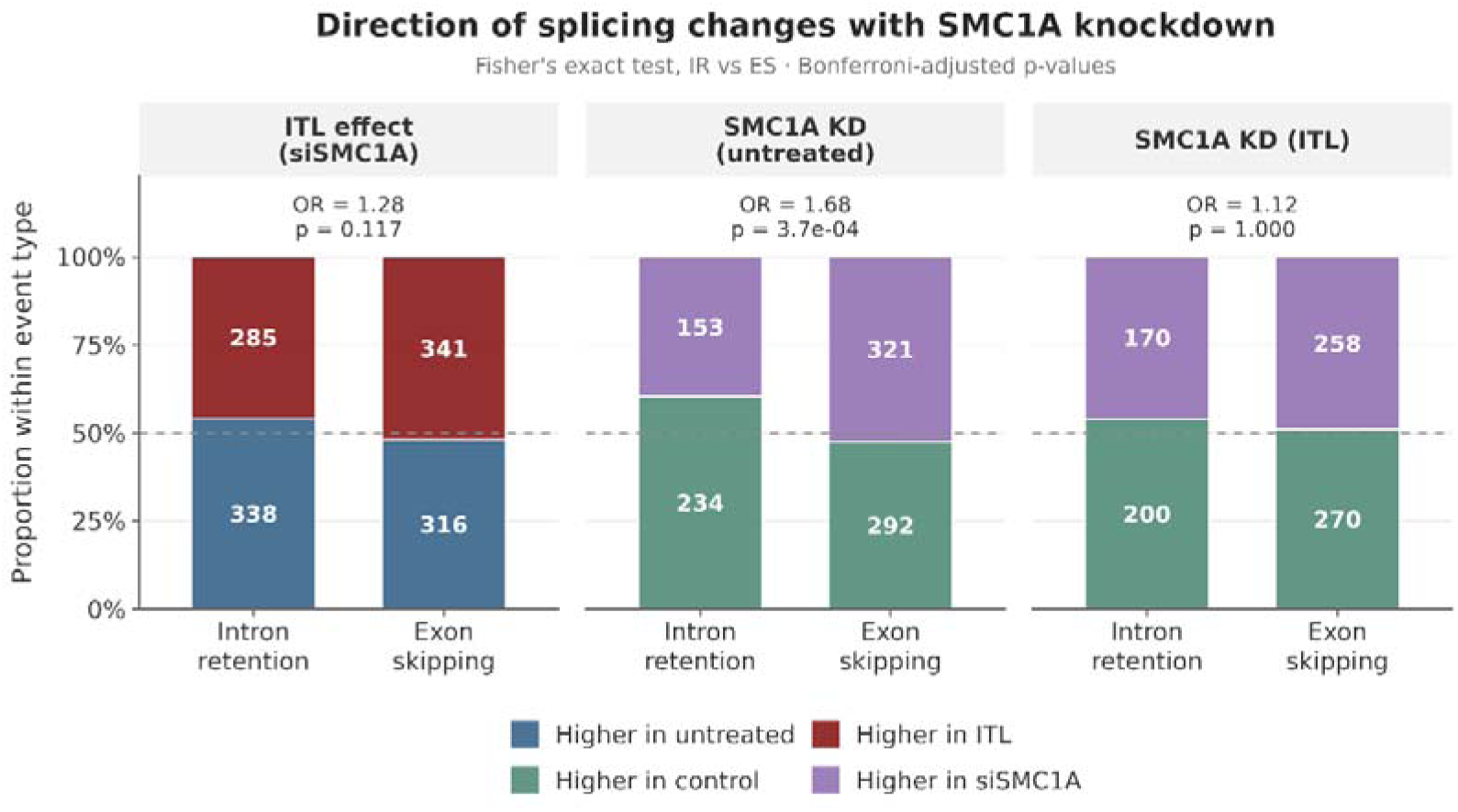
Direction of intron retention (IR) and exon skipping (ES) changes upon ITL stimulation in SMC1a knockdown condition and upon SMC1a knockdown in untreated and ITL conditions. Bars show the proportion of events with higher inclusion in untreated (blue) or ITL-stimulated (red) and in control (green) or siSMC1A (purple) monocytes among events with |ΔPSI|≥0.15; numbers of events are shown inside the bars. Odds ratio and p-value of a Fisher’s exact test comparing the direction of IR and ES events (Bonferroni-adjusted shown here and in Figure 3c).

We then examined the relationship between these changes and nuclear speckle proximity. Introns losing retention upon ITL were enriched among SRSF1-associated RNAs in both control and SMC1A-silenced monocytes, after stratification for baseline retention, whereas introns losing retention upon SMC1A knock-down showed no significant speckle enrichment (Figure 4b). None of the comparisons showed a significant association with LMNA-proximal RNAs.

**Figure 4b.**
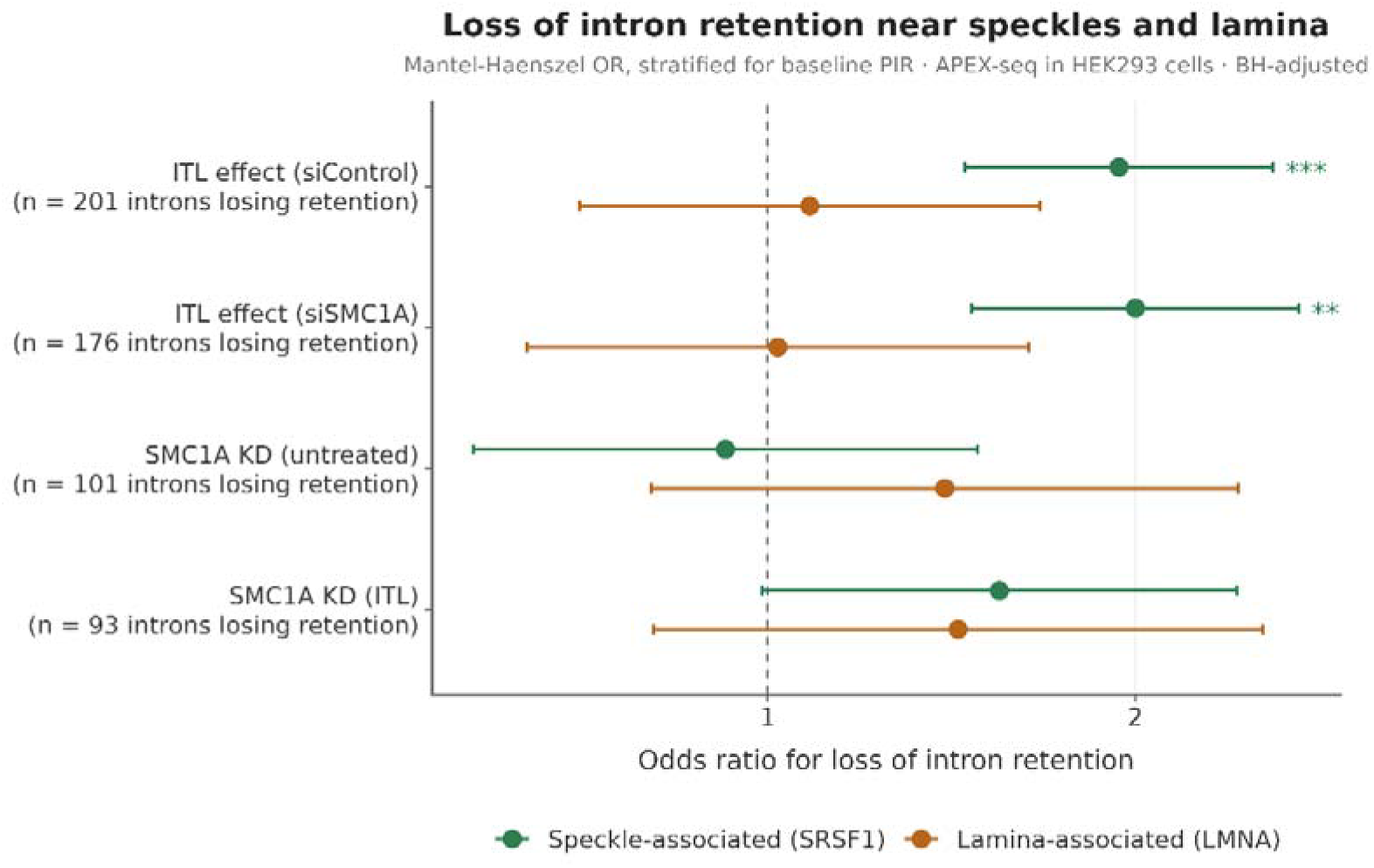
Association between loss of intron retention and nuclear speckle or lamina proximity in four comparisons: ITL versus untreated in control (siControl) and SMC1A-depleted (siSMC1A) cells, and siSMC1A versus siControl in untreated and ITL-stimulated cells. Mantel–Haenszel odds ratios (95% confidence intervals) for an intron to lose retention (ΔPIR≤−0.15, second minus first condition) if it is associated with nuclear speckles (SRSF1, green) or the nuclear lamina (LMNA, orange) in APEX-seq data of HEK293 cells^15^, stratified by quintiles of baseline retention. The number of introns losing retention in each comparison is shown. Only introns quantified in both APEX-seq datasets were included. *, **, ***: BH-adjusted p<0.05, 0.01, 0.001 (eight tests).

We tested whether these splicing changes relate to differential SMC1A binding using ChIP-seq peaks from naïve and stimulated monocytes^33^ in the two comparisons with significant IR/ES imbalance (see above, Figure 3c and 4A). Introns that lost retention upon stimulation were almost 2-fold enriched in SMC1A-depleted genes (log2 enrichment = 0.90, Bonferroni-adjusted p < 0.01), whereas exons with increased skipping upon stimulation were 1.4-fold enriched in SMC1A-enriched genes (log2 enrichment = 0.51, p < 0.01; Figure 4c). This is consistent with loss of SMC1A from speckle-associated regions, since the introns released upon stimulation are speckle-proximal (Figures 3e,f). Of note, IR events in the naïve versus SMC1Akd comparison showed no such enrichment. Thus, the release of speckle-proximal retained introns is a reproducible feature of inflammatory stimulation that occurs in genes losing SMC1A binding and persists at reduced SMC1A levels, whereas SMC1A depletion affects a distinct, non-speckle set of introns.

**Figure 4c.**
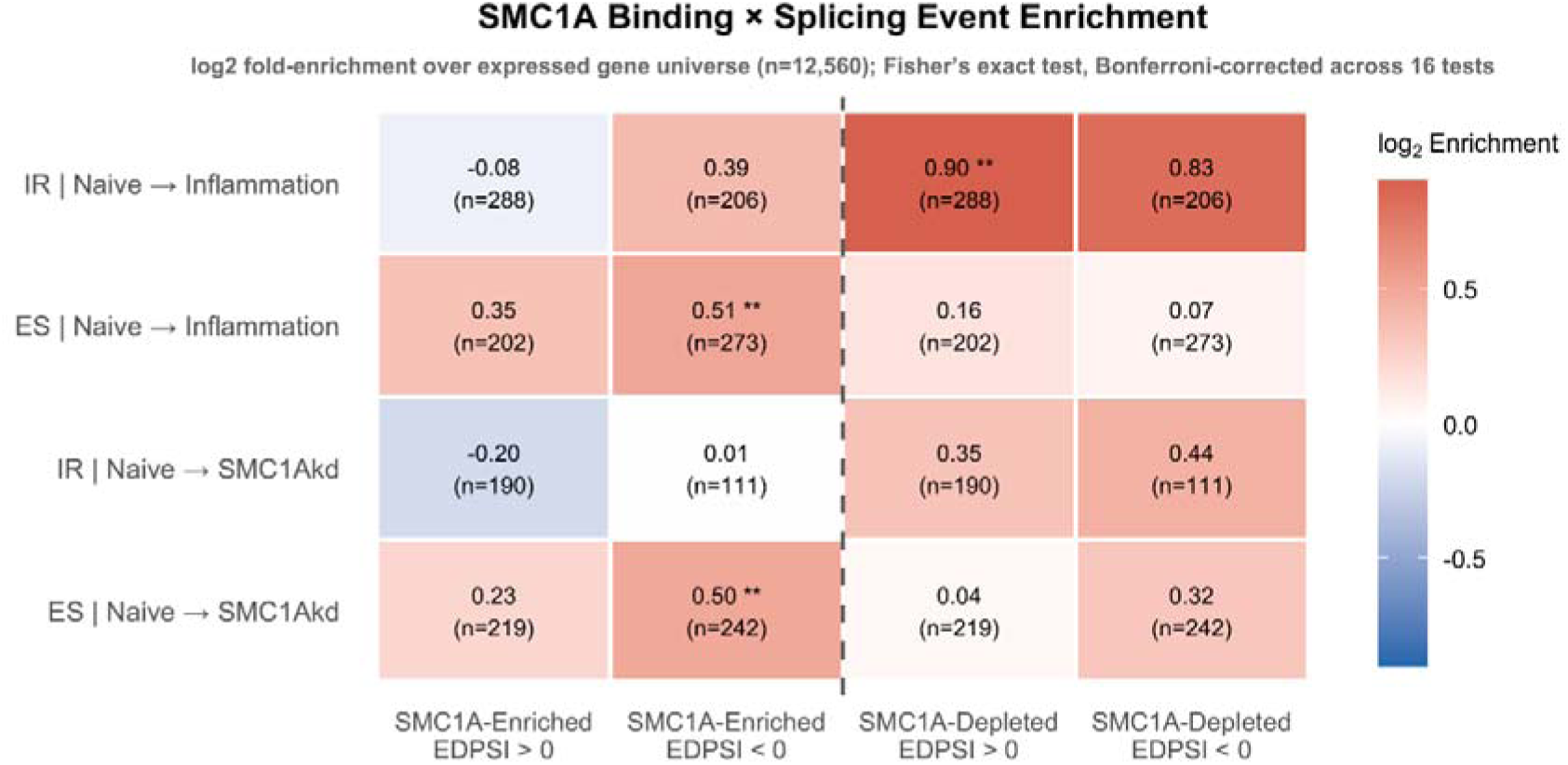
Relative Enrichment calculated as log2-observed/expected ratios of overlaps of SMC1A binding peaks with genes containing IR and ES events in naïve, untreated and ITL-stimulated monocytes. Numbers in parenthesis correspond to numbers of genes in each group. Stars denote significance of a Fisher’s exact test (***<0.001, **<0.01, *<0.05).

### SMC1A is associated with distinct intron subpopulations prior to and post-inflammatory stimulation

Centrally located genes, particularly those associated with NS, tend to have shorter introns with increased GC%^44,46,47^, which promote a splicing mode that proceeds via intron definition and thus makes them more prone for intron retention^16^. A particular case of IR is intron detention, a process whereby the retention of one or more introns leads to the nuclear detention (but not degradation) of the transcript, priming it for eventual splicing and export upon specific signaling cues^48,49^.

Intron detention often occurs in NS-associated, high GC%, short introns with elevated conservation levels and a bias towards the 3’ end of transcripts^50^. Untreated-specific retained introns, which lose retention upon stimulation, were significantly shorter than ITL-specific ones (Figure 3d), whereas no such difference in intron sizes was observed when comparing retained introns in SMC1Akd knock-down conditions, suggesting that the change is specific to stimulation (Supplementary Figure 4a). In line with this, introns retained before stimulation were biased towards the 3\ end of transcripts (Figure 5a), a bias that was not explained by exon number (μUntreated = 7.15, μITL = 7.42, p = 0.47), was absent from exon skipping events and showed no directional preference upon SMC1A knock-down (Supplementary Figure 4b).

**Figure 5a.**
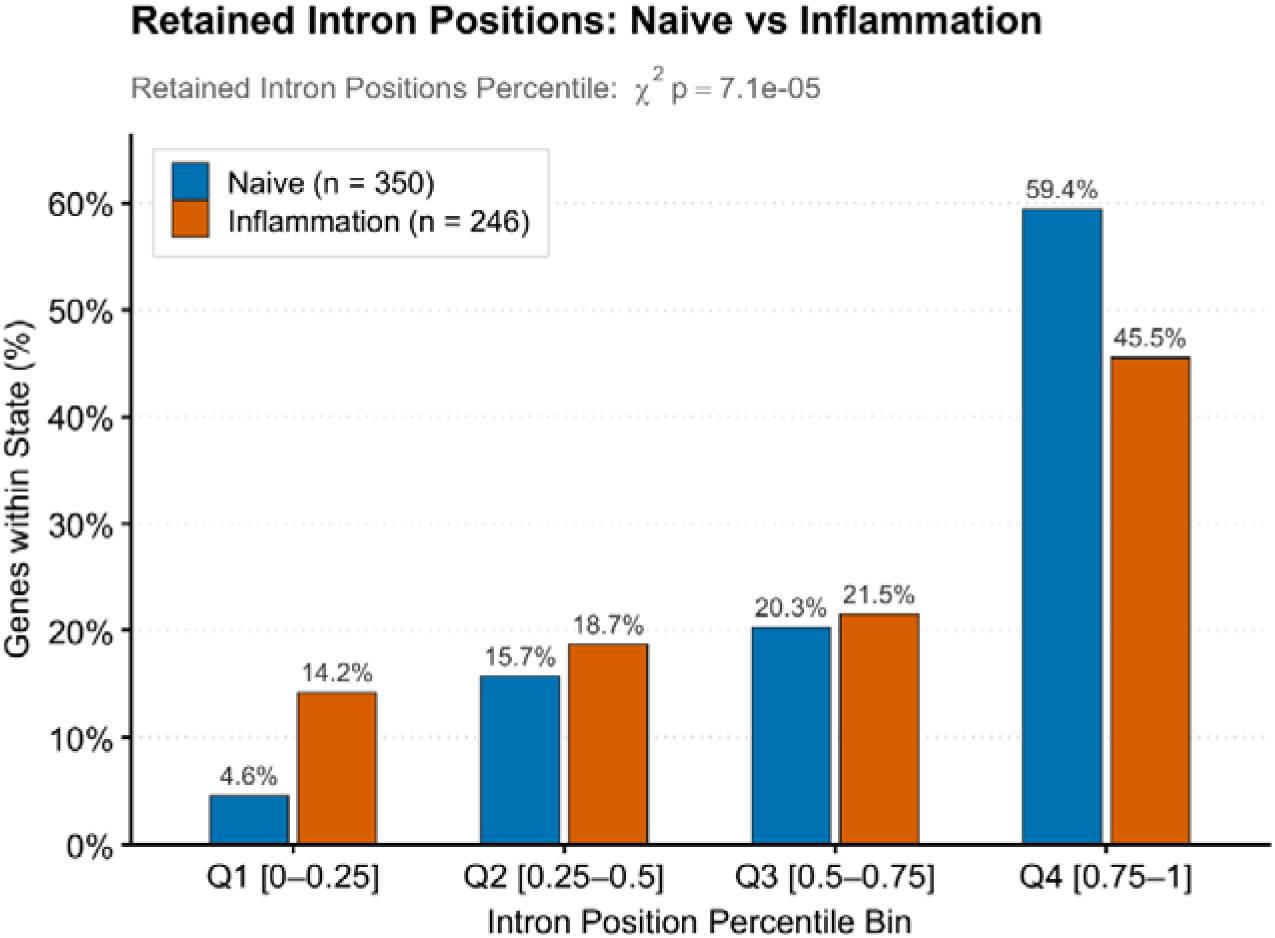
Relative position of retained introns in untreated or stimulated monocytes. Position assigned to relative quintiles corresponding to the position of the intron in the transcript, with 0-0.25 being the most 5’-proximal and 0.75-1 the most 3’-proximal bin (p-value of a χ^2^-test shown in the plot).

Central genes with shorter introns show smaller GC% differences between exons and their flanking introns, whereas this "leveled" architecture yields its place to step-like "differential" GC% patterns in long-intron genes at the nuclear periphery^15,16^. We therefore examined the exon–intron–exon GC% architecture of retained introns within each radial zone. Introns that lost retention upon stimulation had a levelled architecture in every zone, with intron GC% similar to, or higher than that of their flanking exons, often against the general pattern of measured introns from the same zone, which became increasingly differential towards the periphery (Figure 5b).

**Figure 5b.**
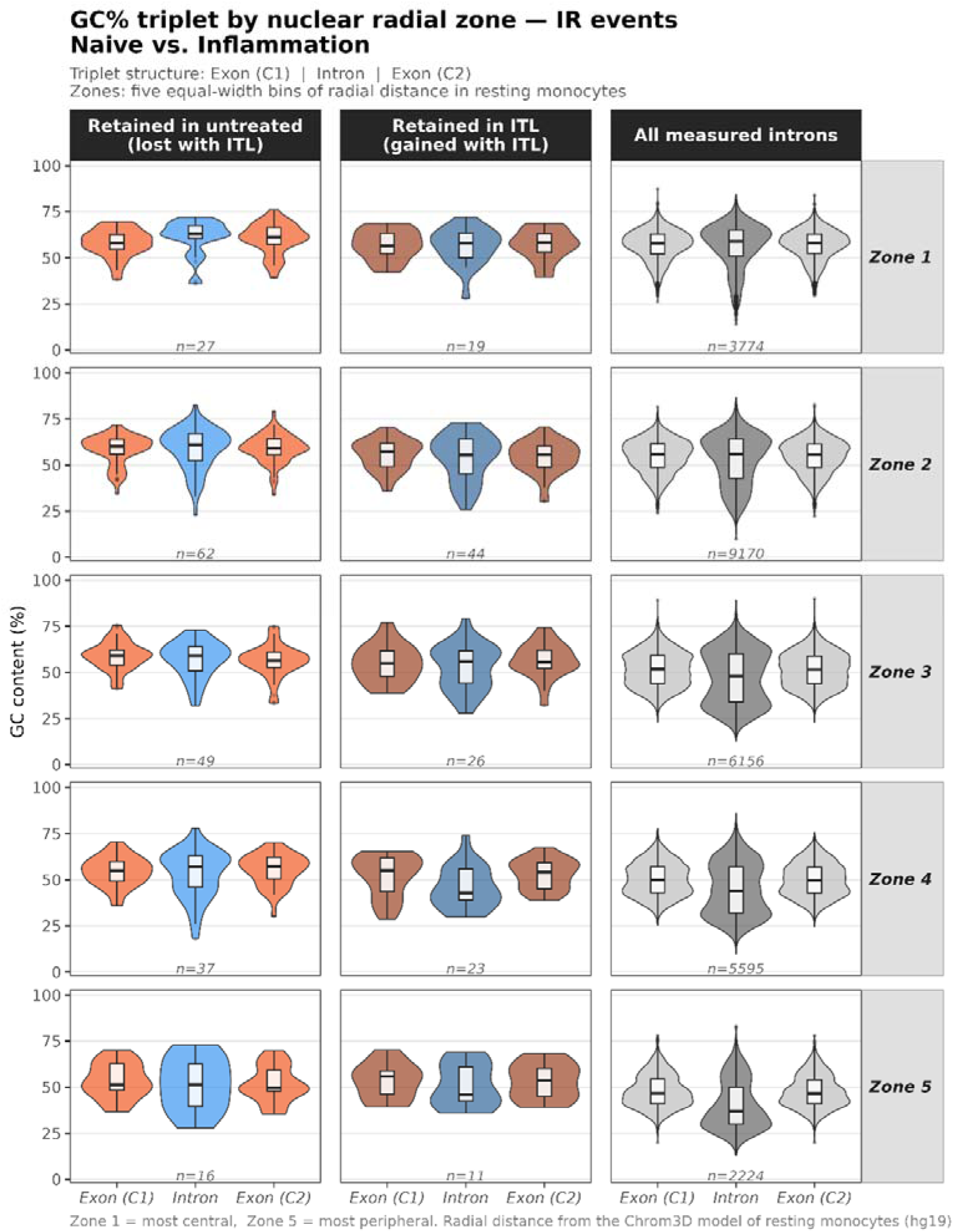
GC% compositional architecture of exon-intron-exon triplets centered around retained introns in untreated (left) and stimulated (center) monocytes compared to total sets of introns (right). Different radial zones defined from our Chrom3D model of untreated monocytes, with increasing radial distance from top to bottom.

Two recent works have highlighted the link between short, high GC% introns and nuclear speckles, going on to propose that protein components of the speckles, such as SON and perhaps even the speckles themselves have evolved to facilitate the efficient splicing of such introns^46,47^. In our data, introns retained before stimulation, i.e. prior to the stimulus-associated loss of SMC1A from speckle-proximal regions, were short and had levelled exon-intron GC% architectures (see Figure 3d, Figure 5B). By contrast, longer introns flanked by shorter exons favour splicing via exon definition, a mode that is correlated with not only faster but also more accurate splicing^51^.

As a proxy of intron versus exon definition splicing mode, we compared the RIME value ratio (Ratio of Intron to Mean Exon, see Methods)^51^ between genes that were SMC1A-enriched or -depleted upon inflammatory stimulation, separately for each of the five radial zones. We found SMC1A-enriched genes to have consistently higher RIME values (Figure 5C), which is compatible with a higher likelihood of splicing through exon definition, independently of their radial distance.

**Figure 5c.**
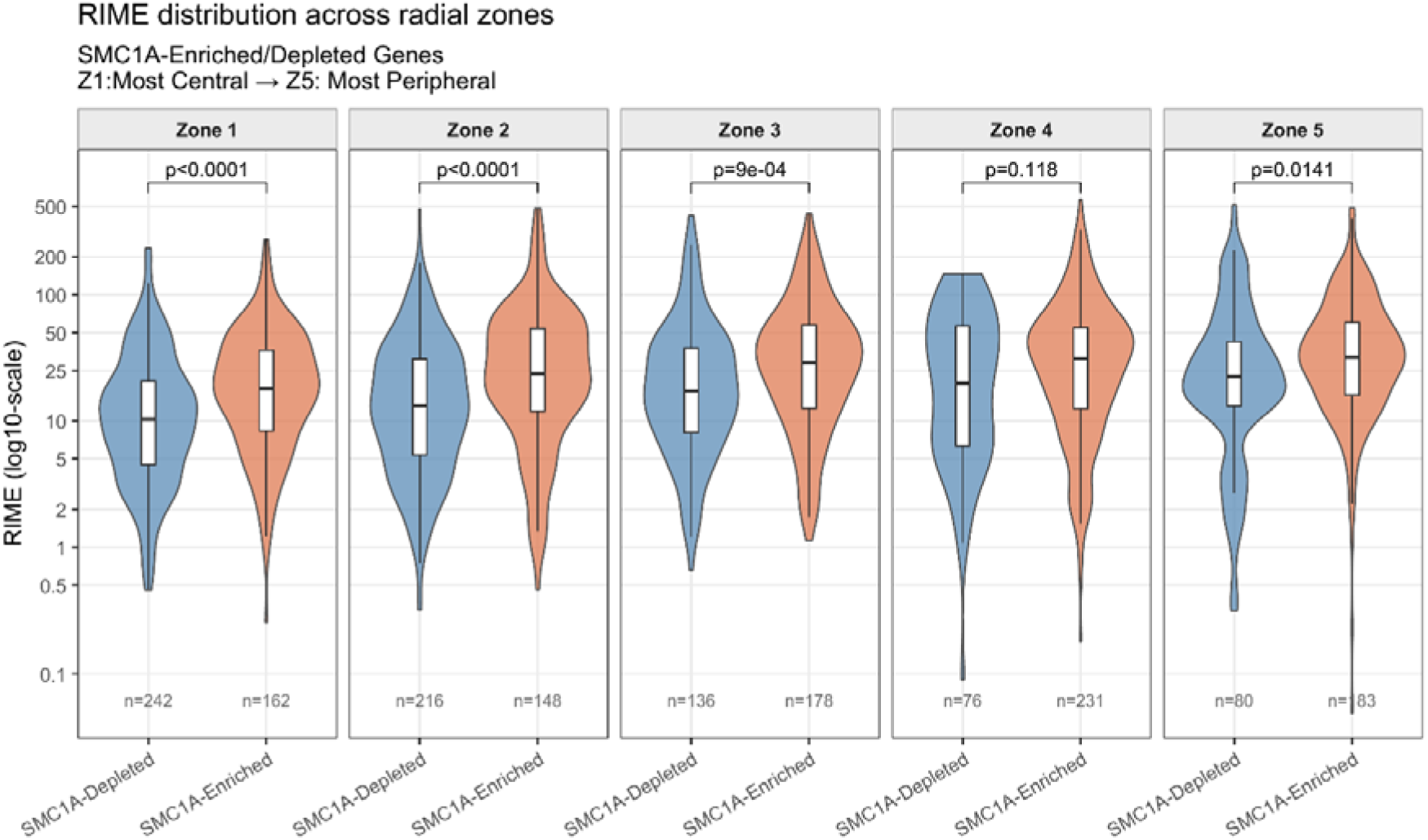
RIME values (as means for all exon-intron-exon triplets) for SMC1A-Enriched and SMC1A-Depleted genes across five radial zones with increasing distance from the genome center. (p-values of pairwise Wilcoxon Rank Sum tests).

### SMC1A-enriched genes show short nuclear mRNA half-lives and exon-intron architectures linked to faster processing

Stress response and inflammatory genes such as cytokines are known to have very fast export times to the cytosol, where they also are relatively short-lived^52^. Particular cases, such as immediate early genes, rank among the most rapidly exported mRNAs. A combination of SLAM-Seq with computational modeling in epithelial cells^53^ showed the nuclear half-lives of immune responsive genes to be among the shortest ones observed. Overall, mRNA stability is largely shaped by the existence of AU-rich elements (AREs) in the 3’UTR and the efficiency of splicing^54^ as early studies have suggested that processed transcripts, which have undergone splicing, are primed for rapid export to the cytoplasm compared to unspliced ones which are processed more slowly^55^.

To relate SMC1A redistribution to mRNA residence times, we compared our gene groups with nuclear and cytosolic mRNA half-lives estimated by SLAM-seq in HeLa-S3 cells (see Methods).

Transcripts of SMC1A-enriched and inflammation-activated genes had short half-lives in both the nucleus and the cytoplasm (Figure 6a). Short cytosolic (but not nuclear) half-lives paralleled enrichment of AU-rich elements (ARE) (Supplementary Figure 5a; Methods), with a significant anti-correlation between ARE enrichment and mean cytosolic half-life (Spearman’s rho=-0.783, p=0.0041).

**Figure 6a.**
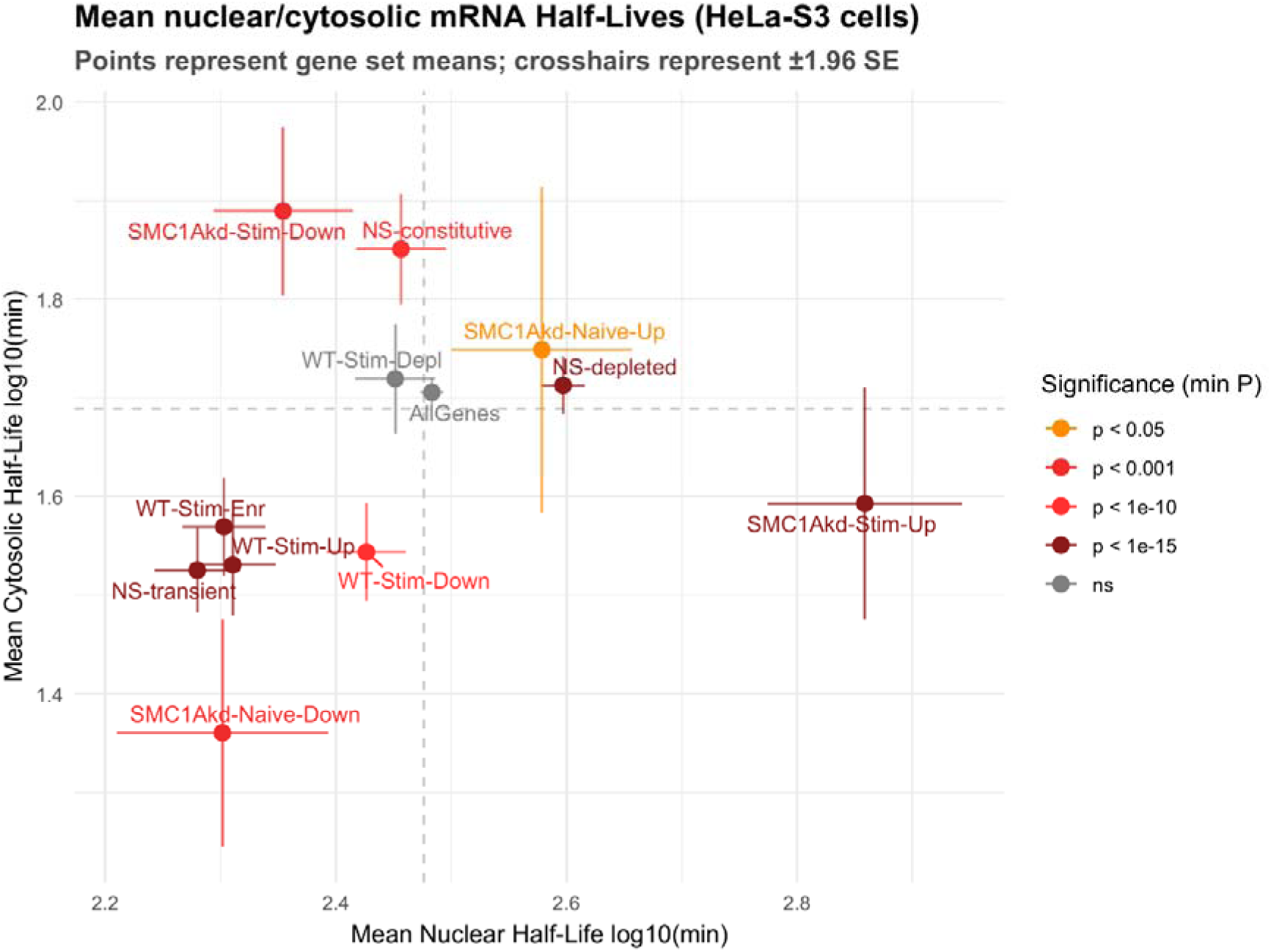
Mean nuclear (x axis) and cytosolic (y axis) mRNA half-lives measured in HeLa cells^53^ for the various gene sets identified in this study. Bars correspond to +/-1.96 standard error. Color coding corresponds to the p-value of a Wilcoxon Rank Sum test against the entire set of genes measured. Grey dashed horizontal and vertical lines correspond to global means for all transcripts.

Contrary to the case of the cytosol, the variation of nuclear half-lives could not be attributed to ARE elements (Spearman’s rho=-0.35, p=0.256). Genes repressed by SMC1Akd in naïve conditions were extremely enriched among the most short-lived mRNAs in both the nuclear and cytosolic compartments (Figure 6a), indicating that SMC1A-dependent genes tend to encode short-lived mRNAs. Under both naïve and inflammatory conditions, transcripts down-regulated by SMC1Akd had significantly shorter nuclear residence times than up-regulated ones (p<10^-5^ and p<10^-16^ respectively).

An identical analysis on data obtained with a different protocol from K562 cells^56^ produced very similar results (Supplementary Figure 5b).

Increasing lines of evidence point out that nuclear residence times are the rate-determining step in overall transcript half-lives^53,56,57^. On the other hand, mRNA processing efficiency has been causally linked with the nuclear export process^58^ as components of the splicing machinery have been shown to colocalize with^59^ and even recruit^60^ protein complexes required for export such as TREX. To test this, we correlated mRNA half-lives (from HeLa-S3 cells) with exon-intron GC% differentials, used as a proxy for exon-definition splicing. Exon-intron GC% architecture was effectively linked with nuclear but not cytosolic half-lives (Figure 6b). Nuclear residence times may therefore be tuned by exon-intron architecture, whereas cytosolic half-lives relate to ARE content; whether this reflects faster processing or export remains to be tested directly. It nevertheless strongly suggests an additional layer of control of mRNA turnover.

**Figure 6b.**
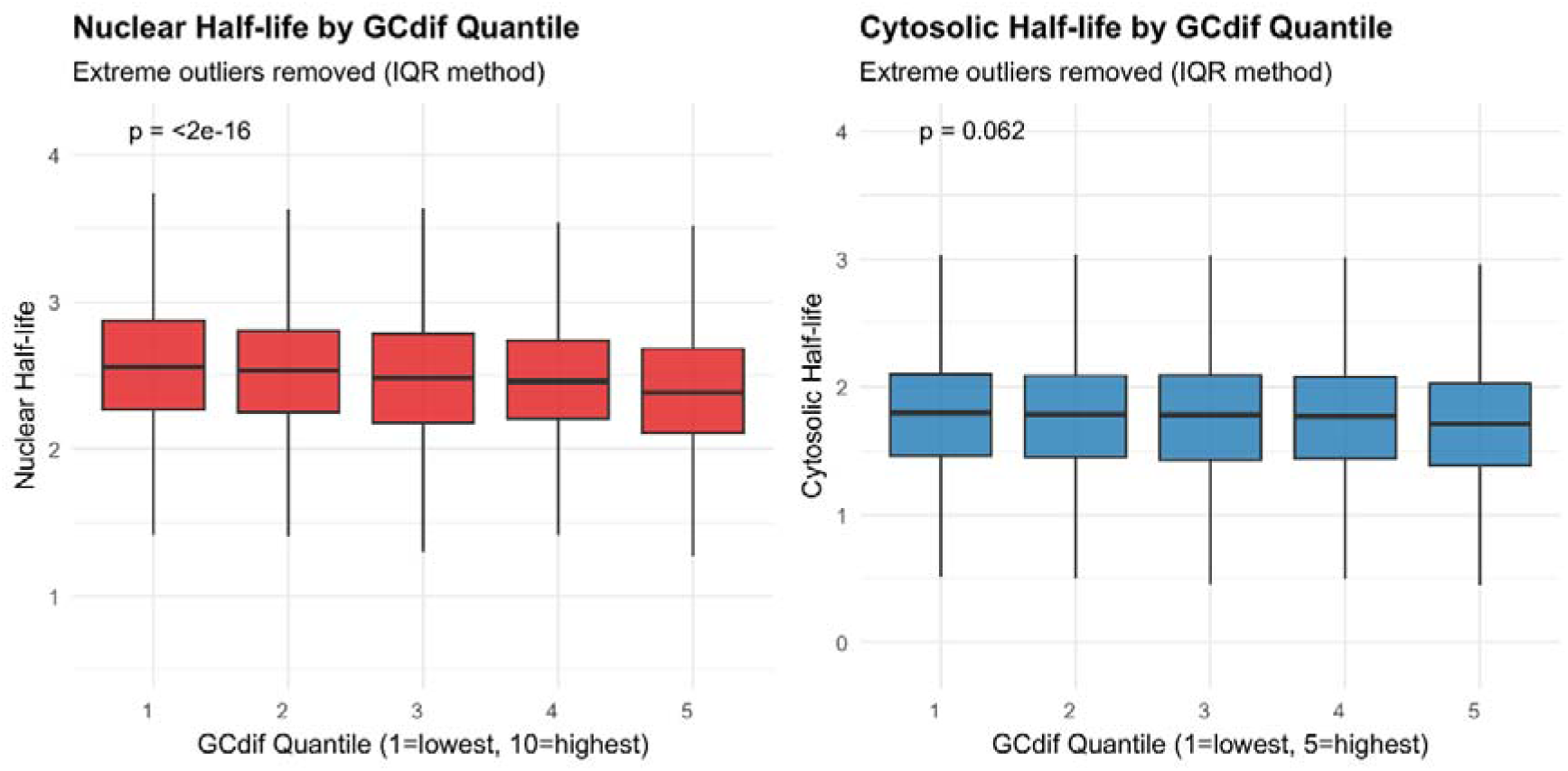
Nuclear (left) and cytosolic (right) half-life times for all transcripts against their mean exon-intron GC% differentials in five quantiles. Half-times estimated for HeLa-S3 cells with SLAM-Seq ^53^(see Methods). GC%-differentials calculated as averages for all internal exons in the same transcript (see Methods). P-values of a Kruskal-Wallis test.

### Radial position and exon-intron architecture covary with gene function

Our analysis connects SMC1A redistribution with gene expression at multiple levels, including transcriptional activation, splicing and, potentially, mRNA processing and export.

While this stresses the importance of nuclear genome organizers in general, it also reflects a functional, underlying genome compartmentalization. We asked whether gene function relates to radial position and exon-intron architecture independently of SMC1A localization. Peripheral enrichment of structurally and transcriptionally more complex genes (greater exon and alternative transcript numbers) and the correlation of exon-intron architecture with higher splicing efficiency are consistent with this^51,61^. A general enrichment of stress-response genes in the nuclear periphery has been shown to be the case in simpler eukaryotes^62–64^ but genes in proximity to the nuclear lamina are also enriched in cell-type specific and developmental functions in the more complex genomes of mammals^34^.

We found strong echoes of a broad, underlying functional organization of the monocyte genome in an SMC1A-independent analysis of gene radial positioning and exon-intron GC% differentials. Using 40 KEGG pathway categories (see Methods), we compared GC% differentials and radial distances against the genome average (Supplementary Figure 5c, see also Methods). We observed a similar trend for basic functions such as transcription and metabolism having low values for both properties. On the other hand, stress-related processes, signaling and secondary metabolic pathways had higher values, suggesting a peripheral positional enrichment with strong patterns of differential exon-intron architecture.

A clear, positive correlation between the two properties was evident at the level of mean aggregates over all genes in each category (Figure 6c). Housekeeping functions related to energy metabolism, protein translation and DNA replication and repair occupied the one end of the spectrum, corresponding to a preference for central positions and levelled exon-intron architectures. Stress-related and cell-type specific functions such as signaling, membrane transport and immune and infectious diseases were clearly located at the other end with peripheral positioning and differential exon-intron structures (Figure 6c). Together, these observations are consistent with gene structure and radial position reflecting gene function.

**Figure 6c.**
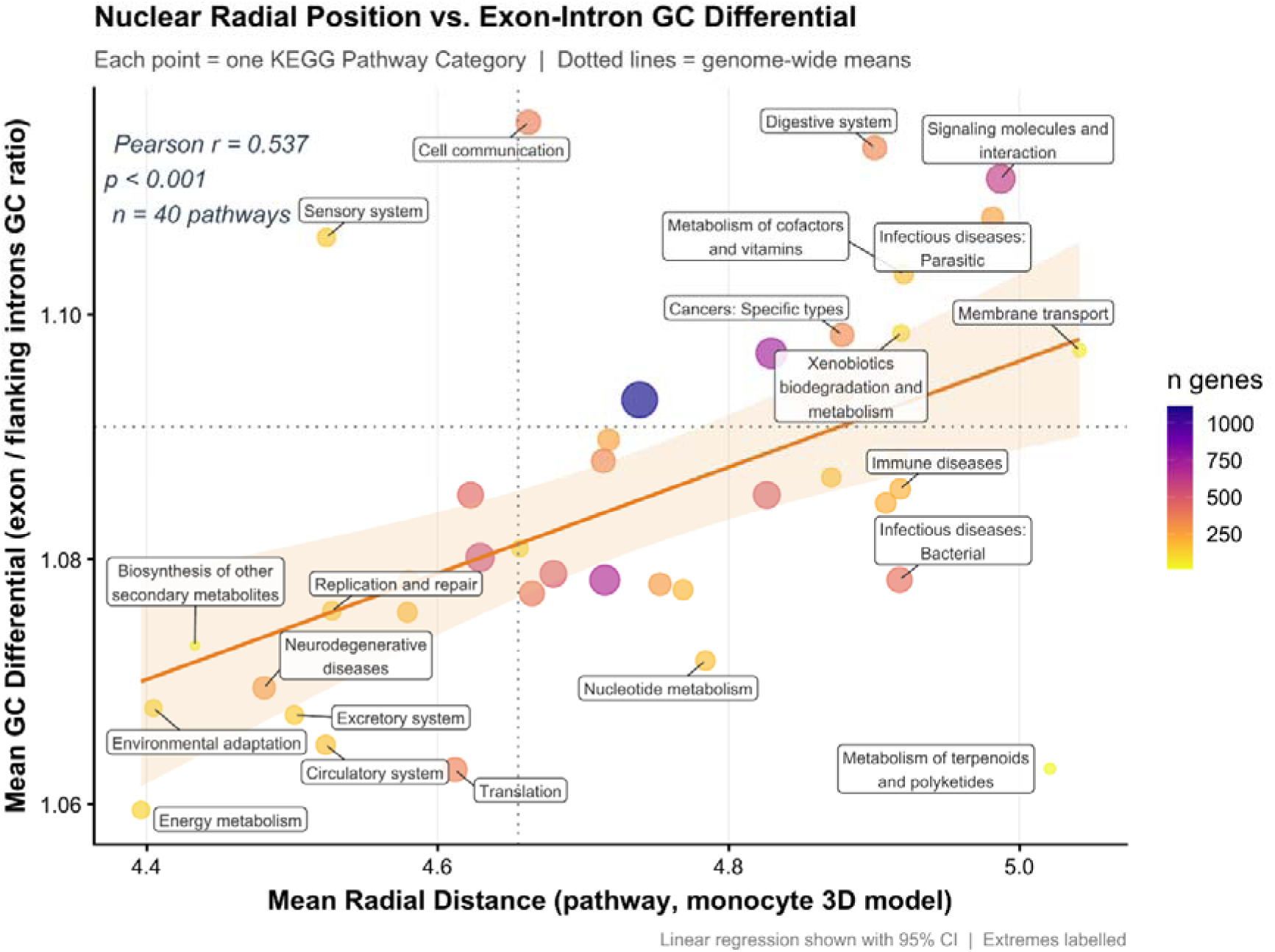
Comparison of mean exon-intron GC% differentials and radial distances obtained from a Chrom3D model of resting monocytes. Each point corresponds to aggregate values over all genes assigned under a KEGG pathway category. Trendline corresponds to linear regression, with coloured band corresponding to 95% confidence interval. Size and color of dots representative of gene set size.

## Discussion

The results we present here, for the behaviour of SMC1A under inflammatory stimulation, align with accumulating evidence regarding the extended roles of nuclear, architectural proteins in various cellular contexts under stress conditions. Our results extend our recent findings^33^ on SMC1A in the transcriptional response to inflammation in human monocytes. Whether this represents a general feature of SMC1A biology across cell types and stress conditions, and whether it extends to the cohesin complex as a whole, remain important open questions.

Our analyses indicate that, upon inflammatory stimulation, SMC1A loses binding at speckle-proximal regions, GC-rich, intron defined genes and gains binding at longer, more peripheral genes, whose splicing likely proceeds through exon definition. These two gene populations differ in composition and architecture within each radial zone, suggesting that SMC1A redistribution follows gene composition rather than merely radial position. The mechanism for this redistribution remains elusive, but a number of hypotheses may be explored. It may involve post-translational modifications that alter SMC1A binding preferences or/and its diffusion rate, or it may proceed through its differential interactions with other, hitherto unidentified protein complexes.

Although specific to one architectural protein and to inflammation, our findings may reflect a more general phenomenon, in which architectural proteins navigate a functionally compartmentalized genome.

Central to this concept is the notion that the overall genome organization, at least in terms of radial positioning, remains generally invariant. In support of this, radial gene coordinates correlated significantly between cell types and methods. A recent work implementing Radial-C, a combination of Mirco-C and MNase gradient diffusion, attempted to capture the changes in radial positioning, brought about the depletion of architectural transcription factors^65^ and found that even upon acute depletion of multiple factors, more than half of the attenuated loops still had the two anchors between very close radial range. Together with the gene properties reported here, this makes extensive genome rearrangements a less likely explanation than a redistribution of SMC1A, although radial positions after stimulation were not measured.

The relationship between SMC1A redistribution and pre-existing chromatin loops provides an additional architectural context for these findings. SMC1A gained sites were enriched at loop anchors located near TAD boundaries, whereas no enrichment was observed at anchors outside these regions. Genes within loops whose anchors gained SMC1A were more frequently induced upon stimulation, particularly when both anchors gained SMC1A. This association may reflect a relationship between SMC1A redistribution and pre-existing chromatin organization. As loops and domains were mapped in resting monocytes, we cannot determine whether loop architecture itself changes upon stimulation. Micro-C analyses in stimulated monocytes could help establish whether these changes in occupancy are accompanied by architectural remodelling and contribute directly to inflammatory gene induction.

A central finding is the loss of SMC1A binding at speckle-proximal regions upon stimulation, which coincided with a reduction in intron retention. The released introns were short, GC-rich, speckle-proximal and biased towards the 3\ end of transcripts, features of detained introns, suggesting that stimulation may release detained transcripts for splicing and export. While this reduction could not be linked to specific functional enrichments, it may be seen as a generalized response that has been already associated with immune cell activation^66^. In various immune cell types, intron retention has been shown to act as a balancing mechanism promoting sustained gene translation^67^, its disruption leading to the activation of splicing factors^67,68^. In this context, reduced intron retention may represent one stage in the remodeling of post-transcriptional processing towards inflammatory gene expression. Importantly, intron release persisted at reduced SMC1A levels, whereas SMC1A knock-down itself affected a distinct, non-speckle set of introns, and the speckle association of SMC1Akd-sensitive genes differed between naïve and stimulated monocytes (Figure 3b). The loss of SMC1A from speckle-proximal genes and the release of their retained introns therefore appear to be coordinated features of the inflammatory response, rather than one being a direct consequence of the other.

Recent works have revealed the role of cohesin in organizing the “ground state” of chromatin around nuclear speckles^25^ and splicing regulation^69^. Our results are compatible with disruption of this state upon extracellular cues: SMC1A binding is lost at speckle-proximal regions upon inflammation and gained at genes whose architecture favours exon-definition splicing, which is faster and more efficient. Genes spliced via exon definition are enriched in the nuclear periphery and carry specific DNA composition patterns^16^. They also carry particular nucleosomal signatures, which, as we have shown in the past, are mechanistically linked to efficient exon definition and subsequent co-transcriptional splicing^70^.

These findings are compatible with a role for SMC1A in transcription and, potentially, in splicing and post-transcriptional processing. Together with the half-life and exon-intron architecture analyses, they point to a functional genome compartmentalization in which DNA composition, gene structure and radial localization co-vary in genes that may be rapidly activated, processed and exported. While SMC1A (and by extension cohesin) is very likely not the only player in orchestrating this, our findings suggest that nuclear, architectural proteins may mediate complex regulatory outcomes by navigating a pre-existing, underlying genome compartmentalization.

Our work also offers a novel view on how such a functional compartmentalization of the eukaryotic genome may have been shaped to enable such responses. Through a combination of functional enrichment and gene structure profiling, we show how genes with constitutive functions and genes associated with stress responses differed in radial position and exon-intron architecture (Figure 6c), a layout that could allow stress-response genes to remain poised for rapid activation while constitutive genes maintain steady expression. The stress-induced redistribution of nuclear proteins with multifaceted regulatory roles from the center to the periphery, such as the one we describe here for SMC1A, would shift the balance towards stress-response genes without severely disrupting constitutive ones, by effectively deploying the genome structure as their substrate.

Our findings suggest several testable predictions. First, radial positions and chromatin loops may be mapped in stimulated monocytes, together with direct imaging of SMC1A relative to nuclear speckles, to test whether the redistribution inferred here occurs in situ. Second, if loss of SMC1A from speckle-proximal regions contributes to intron release, its magnitude should correlate quantitatively with the extent of intron retention change across genes. Third, the contribution of radial position can be tested by relocating selected SMC1A-responsive loci with CRISPR-based tethering and, to separate position from sequence architecture, by editing their intron length and GC% in situ without changing their position.

## Methods

### Experimental setup and comparisons between conditions

This study utilized integrative analysis of gene expression and binding datasets derived from primary human monocytes. Monocyte isolation and culture, ITL doses/duration, siRNA design and knockdown efficiency, RNA-seq and ChIP-seq library preparation and sequencing, donor and replicate numbers, QC are described in our previous work^33^.

RNA-seq experiments were performed under basal and lupus-like inflammatory conditions that included the combined induction with IFN, TNF and LPS (ITL stimulus), to identify transcriptional changes. Genes significantly upregulated or downregulated upon inflammatory stimulation were classified as “Stimulation Activated (Up)” and “Stimulation Repressed (Down)” respectively. In parallel, ChIP-seq targeting SMC1A was conducted to determine changes in chromatin binding upon lupus-like inflammation. Genes exhibiting increased or decreased SMC1A occupancy were designated as “SMC1A - Enriched” and “SMC1A - Depleted”. As all experiments involved wild-type monocytes, gene sets were given the prefix (WT).

Additionally, RNA-seq following SMC1A knockdown (SMC1Akd) was performed under both untreated and inflammatory conditions. This analysis yielded two further gene categories: “SMC1Akd-Naive” Activated or Repressed (genes affected by knockdown under basal conditions) and “SMC1Akd-Stimulated” Activated or Repressed (genes differentially expressed only under inflammatory conditions with reduced SMC1A levels). All RNA-seq and ChIP-seq datasets were generated from primary human monocyte cultures and are available in GEO under accession number <u>GSE279726</u>.

Gene lists are provided as Supplementary File 1 (see Supplementary Code and Data), while differentially enriched and depleted SMC1A binding sites (Figure 1C) are provided in https://github.com/christoforos-nikolaou/SMC1A-in-Inflammation.

### Gene Features and Machine Learning

We compiled compositional, sequence constraint, structural and transcriptional complexity properties for all human genes in the following way. Gene annotations for the human genome (GRCh38/hg38) were obtained from GENCODE. Gene length, distance from the closest gene, number of associated transcripts and maximum exon count (over all transcripts) were obtained directly from the gtf annotation file. GC% was directly calculated for gene, exon and intron coordinates provided by Ensembl. Mean exon and intron GC% values were calculated for all exon and introns assigned to each ENSEMBL Gene ID (see below for exon, intron analysis in more detail). Sequence conservation was calculated for the exact same coordinates based on aggregate phastCons values, obtained for hg38. Alternative splicing events (as ES: Exon Skipping, IR: Intron Retention and Alt-DA: Alternative Donor/Acceptor Sites) were obtained from VAST-DB^71^ (Supplementary File 2).

We performed a Random Forest (RF) Classification on the entire set of genes for which we could compile data (n=19912). This was followed by a SHAP (SHapley Additive exPlanations) analysis in model’s predictions to assess the exact contribution of each feature. SHAP analysis assigns a numeric value to every feature and for every instance (gene) in the model, in order to explain whether it pushed the prediction in the multiclass classification model higher or lower. RF and SHAP analyses were implemented in R with custom scripts provided as Supplementary Code. SHAP values are provided as Supplementary File 3 (see Supplementary Code and Data).

### Interaction Analysis

Starting from 12,833 expressed genes (CPM ≥ 1 in at least one condition), a Welch t-test was run independently for each of the three main contrasts against CU: siRNA (SU vs CU), ITL (CITL vs CU), and the combined treatment (SITL vs CU). BH-FDR correction was applied within each contrast. The gene set for clustering was defined as the union of all genes reaching FDR < 0.05 in any one of the three contrasts, with no LFC threshold applied. This yielded 2,716 genes, each of which had three measured vectors relative to the control/untreated state: A = ITL effect (log2FC of ILT vs Untreated), B = SMC1Akd effect (log2FC of SMC1Akd vs Untreated) and AB = SMC1Akd+ITL effect (log2FC of SMC1Akd+ITL vs Untreated). Interaction was then calculated as I = AB − (A+B) to assess the deviation from perfect additivity.

Genes were subsequently clustered according to the four values (A, B, AB and I). Before clustering, the B and I dimensions were up-weighted by a factor of 2 relative to A and AB — the rationale being that the SMC1Akd and interaction effects are small in absolute magnitude and would otherwise be dominated by the large ITL signal. The four weighted features were then z-scored to unit variance. Ward-linkage hierarchical clustering was applied on Euclidean distances in this standardised space. A choice of k = 6 was chosen after visual inspection of the data. Clustered data are provided as Supplementary File 4 (see Supplementary Code and Data).

### Modeling of the 3D genome and radial positioning

A CD14+ monocyte-specific 3D genome model was generated with Chrom3D^36^ using Hi-C data derived from primary human monocytes of a healthy male donor, that can be found in ENCODE with Experiment Code: <u>ENCSR236EYO</u>. Initial analysis of the Hi-C maps was performed with HiC-Pro^38^. The resulting TAD assignment was extended to allow for a uniform, gap-free tiling of chr1–chr22 and chrX into uniquely labelled beads suitable for use as the input “bead” file in Chrom3D modelling. In accordance with the Chrom3D pipeline recommendations, intra-chromosomal interactions were extracted at 50 kilobase (kb) resolution using the Juicer Tools software^72^, with the binning performed on fixed base-pair coordinates and the resolution explicitly set to 50kb. Inter-chromosomal interactions were created with a similar approach at 1Mb resolution maps of each chromosomal pair. Lamina-associated domains were incorporated from (GSE109924) and lifted-over to hg38. Chrom3D beads were defined as TAD-bounded genomic segments encoded in a GTrack file and then modeled with Chrom3D in a cmm file (Supplementary File 5), which was visualized with ChimeraX^37^ (see Figure 2A).

Radial positioning was calculated in the form of Euclidian distances of each bead (TAD) from the center of the model (x=0, y=0, z=0). Each gene was assigned the radial distance of the bead it was contained in. Euclidian distances were binned into five zones in a way similar to the one described in^16^ with Zone 1 corresponding to the most central and Zone 5 the most peripheral regions (Supplementary File 6, see Supplementary Code and Data).

Analysis of Radial Zone enrichment was performed as observed over expected ratios of genes belonging to each Zone, assessed with a two-sided Fisher’s test.

### Chromatin loop and TAD boundary analyses

Chromatin loops and topologically associating domains (TADs) were obtained from ENCODE Hi-C data identified in resting human CD14+ monocytes (ENCSR236EYO). We used 24,471 loops (ENCFF214LRJ) and 2,576 domains (ENCFF992XHP), both originally mapped to GRCh38. Loop anchors were converted separately to hg19 using UCSC liftOver, while preserving their loop identifiers. We retained only intrachromosomal loops for which both anchors mapped uniquely to the same chromosome and maintained their original order, resulting in 24,382 loops. TAD boundaries were defined as the start and end positions of each domain and were similarly converted to hg19.

Differential SMC1A sites were obtained from the DiffBind analysis of ITL-stimulated versus untreated monocytes and classified as gained (n = 6,839), lost (n = 1,131) or unchanged (n = 17,593), using a p-value threshold of 0.01 and fold-change cutoffs of ≥1 and ≤−1 for gained and lost sites, respectively. Sites were considered associated with loop anchors if their genomic coordinates overlapped an anchor, and with TAD boundaries if their midpoint was within 10 kb of a domain border. Based on these criteria, we classified sites into four groups: overlapping both an anchor and a TAD boundary, overlapping an anchor only, associated with a TAD boundary only, or neither. Enrichment of gained and lost sites in each group was assessed against unchanged sites using Fisher’s exact test, with Benjamini-Hochberg correction for eight comparisons.

We next examined the relationship between SMC1A binding at loop anchors and inflammatory gene expression. For this analysis, we used 8,664 genes with available DESeq2 results for ITL versus untreated monocytes and genomic coordinates in hg19. Genes were assigned to loops when their entire gene body was located between the two anchors. Loops were classified according to whether one or both anchors gained SMC1A, whether they contained SMC1A losses only, or whether their anchors showed no differential binding. For genes contained within multiple loops, we prioritized loops with gains at both anchors, followed by gains at one anchor, loops with losses only and loops without changes. Genes outside all loops were considered separately. Induced and repressed genes were defined by an adjusted p-value <0.05 and a log2 fold change >1 or <−1, respectively. The frequency of induction and repression in each loop category was compared against genes in loops without differential SMC1A binding using Fisher’s exact test.

### Differential Splicing

Alternative Splicing (AS) events were identified utilizing Vast-tools (VAST-TOOLS v2.5.1), the Vertebrate Alternative Splicing and Transcription Tools^71^. Reads were aligned to the human reference genome (hg19), and unmapped reads were subsequently aligned to a comprehensive library of predefined splice junctions. Specifically, four different, dual comparisons were performed to assess differential splicing patterns between conditions in human monocytes. Percent spliced-in (PSI) values were computed for each event, and events with significant changes were identified based on a minimum ΔPSI threshold (ΔPSI, threshold>0.15). The analysis focused primarily on exon skipping (ES) and intron retention (IR) events. All significant ES and IR events for all comparisons are provided as Supplementary File 7 (see Supplementary Code and Data).

### Direction of splicing changes

Events with |ΔPSI| ≥ 0.15 were classified according to the direction of their inclusion changes. In the VAST-TOOLS diff output used here, E[dPsi] represents the difference between the first and second conditions. Thus, for the untreated versus ITL comparison, positive values indicate higher inclusion in untreated cells and, for IR events, reduced retention upon stimulation. The direction of IR and ES changes was compared using Fisher’s exact tests, with Bonferroni correction across the four comparisons (ITL versus untreated in siControl and siSMC1A cells, and siSMC1A versus siControl in untreated and ITL-stimulated cells).

### Intron architecture

Introns quantified in the untreated versus ITL comparison in siControl cells were classified as losing (ΔPIR ≤ −0.15) or gaining retention (ΔPIR ≥ 0.15), with ΔPIR defined as PIR in ITL-stimulated minus untreated cells. All remaining measured introns were included in the comparison group. Intron coordinates (hg19) were obtained from the VAST-TOOLS inclusion table, and intron length was calculated from these coordinates. Intron and flanking exon GC% values were obtained from Tammer et al, by matching intron boundaries to the coordinates of the adjacent exons. Exon–intron GC% difference was calculated as the mean GC% of the two flanking exons, or of the single matched exon when only one was available, minus intron GC%. Groups were compared using two-sided Wilcoxon rank-sum tests, with Benjamini–Hochberg correction across nine comparisons (three variables and three pairwise group comparisons).

### GC% differential and RIME analysis

We obtained the complete set of gene annotation for hg19 in gtf format from Gencode (https://www.gencodegenes.org/human/release_19.html), from which, we extracted all exon and intron coordinates. Intron coordinates were calculated with a for-loop that scans each row of the dataset, utilizing exon coordinates for transcripts with more than one exon. In brief, for each gene or transcript entry, intron start and stop fields were initialized. When an exon was encountered, the loop calculated intron boundaries based on the end of the previous exon and the start of the current one, also taking into account the strand orientation. The resulting intron positions were stored in newly created “intronstart” and “intronstop” columns.

Following coordinate assignment, additional intron-level metrics were calculated. The length of each intron was determined from its genomic start and stop positions. Each intron was also assigned an ordinal number based on its order within the transcript (e.g., first intron, second intron, etc.). These values were stored in new columns and used in subsequent analyses for introns of interest following the results of the differential splicing analysis.

GC% was calculated with an in-house script utilizing FASTA sequences of genes, exons or introns. We then went on to report a triplet of GC% values for every internally annotated exon in the format of intron-exon-intron. For the purposes of analyzing our retained introns, we reversed this order by placing the intron in the middle and flanking it with the previous/next exon in every case. We performed the same analysis for a set of random exons and introns. The file containing the GC% differentials for our IR events and the random exons/introns is provided as Supplementary File 8 (see Supplementary Code and Data).

RIME values were calculated as ratios of the intron length over the arithmetic mean of the two flanking exons as described in^51^. All RIME values alongside the complete set of introns for all transcripts are provide as Supplementary File 9 (see Supplementary Code and Data).

### mRNA half-life analysis

Estimates of Nuclear and Cytosolic mRNA half-lives through modeling of SLAM-Seq data in HeLa-S3 were obtained from a recent work^53^, provided as supplementary data by the authors. Extreme values (top 0.1%) were trimmed from the dataset as outliers and remaining values were then mapped to our list of genes and then mapped on our genes of interest. Mean values and standard deviations were calculated with custom R code (see Supplementary Code and Data).

RNA-flow data were obtained for K562. RNAflow provides a quantitative flux estimate for subcellular residence times, combining TimeLapse-Seq data coupled with GRAND-SLAM and Bayesian ODE modeling. We obtained mean estimates for half lives for nucleus, chromatin, cytosol and polysome, alongside nuclear export rate as provided in the supplementary data of the original work^56^. Replicate 2 (K562_rep2) was used since data mapped on a slightly larger number of our genes of interest. (Replicates 1 and 2 had a mean Pearson correlation of ∼0.9)

ARE data were compiled from ARED-Plus^73^. Entries matching transcript IDs were merged and the final analysis was performed on a binary outcome of any given gene containing a documented ARE. Enrichments were calculated on the basis of a chi-square test (see Supplementary Code and Data).

### Functional Gene Annotation

We used gProfiler^74^ for the functional enrichment analysis and for all supplementary analysis where over-representation enrichment was required. For the analysis presented in Supplementary Figure 5c and Figure 6c KEGG annotation was used. This was implemented in the following way. All KEGG pathways were retrieved from (https://www.genome.jp/kegg/pathway.html) and all unique genes belonging to pathways under the same broad category (e.g. Immune Systems, Cell Motility, Membrane Transport etc) were grouped under a single term. Pathways related to Category 7: Drug Development, were not included (Supplementary File 10).

All genes from each category were mapped on our Radial Distance Data (Supplementary File 5) and GC% differential calculations (Supplementary File 9) and compared to the overall mean with pairwise Wilcoxon Rank Sum tests. P-values were adjusted with Benjamini-Hochberg. Mean values for GC% differential and Radial Distances were obtained for each category and used in a linear regression model. Outliers were detected as top/bottom quintiles (20%) according to their residual values (see Figure 6c).

### External Datasets

GPSeq^5^ data were obtained from GEO under Accession Number: <u>GSE135882</u> for HAP1 and GM06990 cells. Scores were transformed to bigwig format and aggregated over genomic intervals corresponding to the TADs defined in our monocyte cells to allow for a one-to-one comparison with our radial distance. Both distance values and GPSeq aggregate scores were scaled to a 0-1 range for better representation purposes.

Radial distances and radial zone assignments for >10000 genes in K562 cells were obtained directly from the original work^16^ as provided in their Supplementary Data (Table 4). TSA-seq data from K562 cells were obtained from the same supplementary dataset, in the form of proximity percentiles. Genomic regions were assigned a TSA-Seq percentile score based on their proximity to NS, with 100% indicating the closest proximity and 0% the greatest distance.

Nuclear Speckle associated gene sets of three different types (Constitutive, Transient, Depleted) were obtained directly from a recent work^44^ as provided in Supplementary Table 1.

SPIN data for K562 cells, were obtained as supplementary data from the original publication^45^ as genomic coordinate file (bed) for hg38. SPIN uses a Hidden Markov Model to identify genomic localization proximity to distinct nuclear structures, primarily positioning genes on Nuclear Speckle – Nuclear Lamina axis. Enrichment analysis was performed directly on the coordinate data against the genomic coordinates of the studied genes (lifted over to hg38) using custom scripts based on bedtools^75^. Coordinate intersection was used to calculate observed overlaps and these were compared to expected ones under a null model of independence. Significance was assessed on the basis of a permutation test, which included a genome reshuffling of the SPIN states maintaining the number of segments and size distribution for each state (N=1000).

APEX-Seq data on SRSF1 and LMNA-associated intron retention (IR) events were obtained in the form of Supplementary Data from the original publication^15^. Common IR events with our own (|ΔPSI|>0.15) were retrieved. Correlation coefficients of ΔPSI values were calculated as Spearman ρ.

### Statistics and reproducibility

Statistical tests are specified in the figure legends and the relevant Methods sections. They comprised Wilcoxon rank-sum, Welch t, Fisher’s exact, chi-squared, Kolmogorov-Smirnov and Kruskal-Wallis tests, Pearson and Spearman correlations, and permutation tests, with Benjamini-Hochberg correction where indicated.

### Data and code availability

Supplementary Data from this work have been deposited to Zenodo (https://doi.org/10.5281/zenodo.18266710). Extended methods and original code written for the presented analyses may be found in the accompanying github site (https://github.com/christoforos-nikolaou/SMC1A-Genome-Architecture), alongside external datasets (see above) and extended code, for the production of all Figures.

## Acknowledgements

We thank all members of the Nikolaou and Papantonis labs for valuable discussions related to the findings and the analyses presented in this work.

## Author contributions

CN, SP and MM conceived of the study, compiled data, performed analyses and interpretation of the data. AK, AS, DT and CM performed bioinformatics analysis. GB, DK, AC provided initial data and performed experiments. AP and AM performed experiments. GB secured samples and funding. AP, GB and CN supervised experiments and analysis. CN and SP wrote the manuscript. GB and AP provided expert opinion and corrected drafts of the manuscript. All authors read and approved the final version of the manuscript.

## Supplementary Information

## Supplementary Information

**Supplementary Figure 1a.**
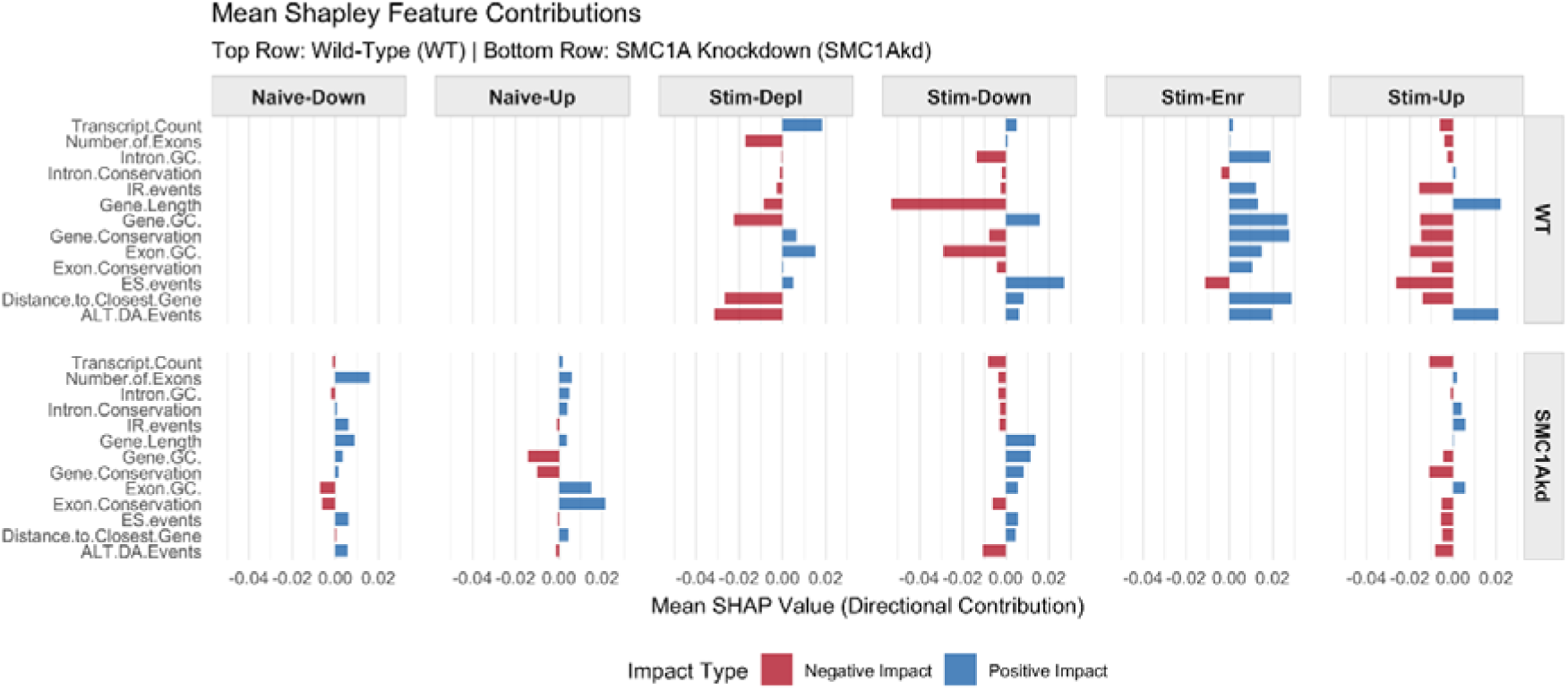
SHAP values obtained from a multiclass classification Random Forest Model. SHAP impact values evaluate the per-class probability for each gene to belong to the given class compared to all other classes.

**Supplementary Figure 1b.**
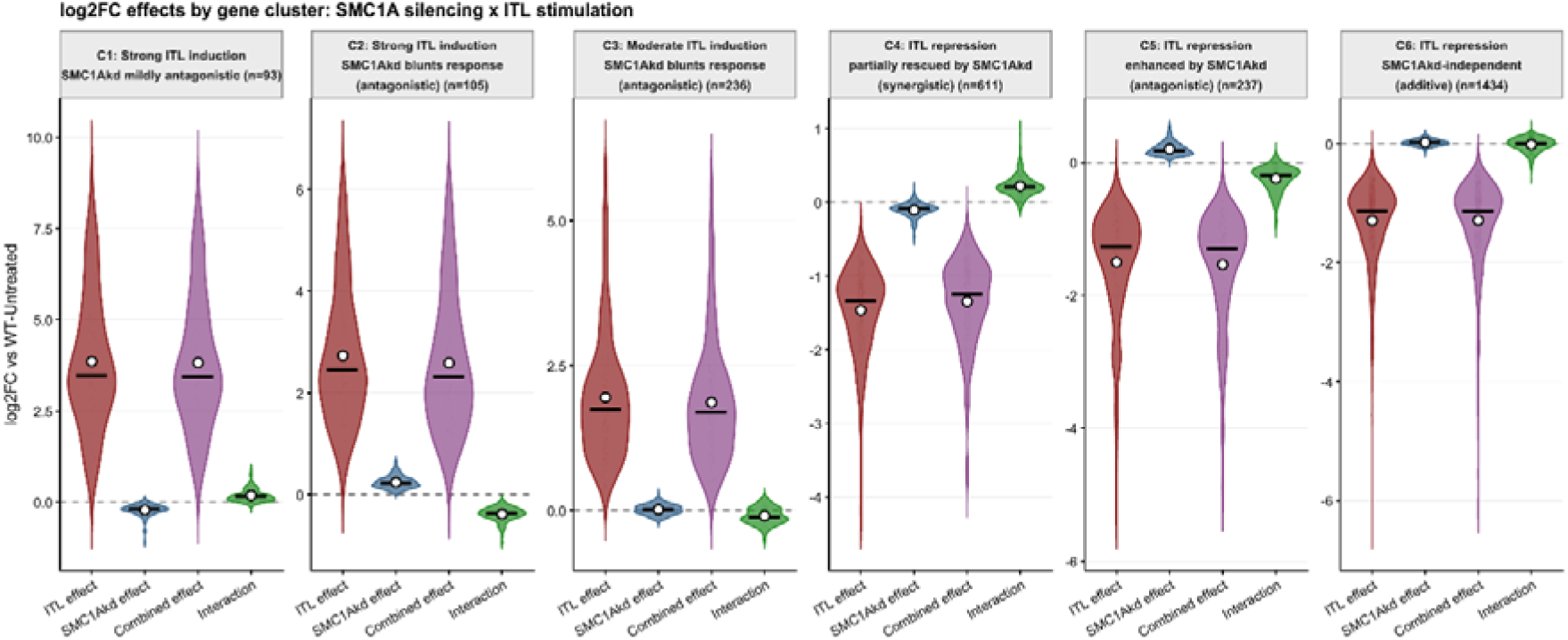
Interaction Analysis of Gene Expression Changes for Inflammatory Stimulation (ITL, red), SMC1Akd (blue) and their combination (purple). Interaction (green) is measured as the log2fold-change of the combination minus the sum of the two independent effects (I = AB – (A+B), see Methods).

**Supplementary Figure 1c.**
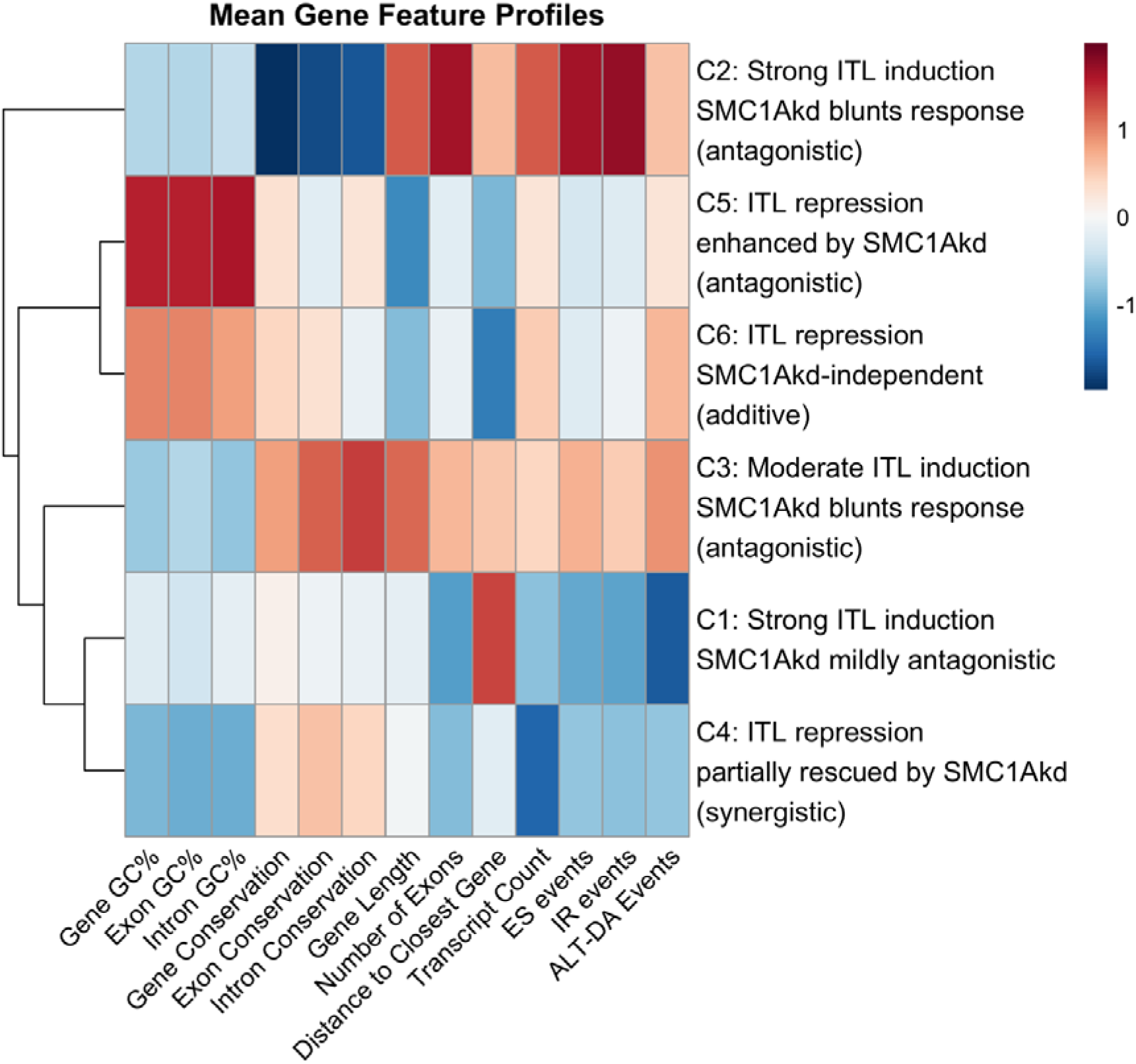
Average Gene Profile Analysis (as in Figure 1d) for the six gene clusters deduced in the Interaction Analysis (see Methods and Supplementary Figure 1b)

**Supplementary Figure 2.**
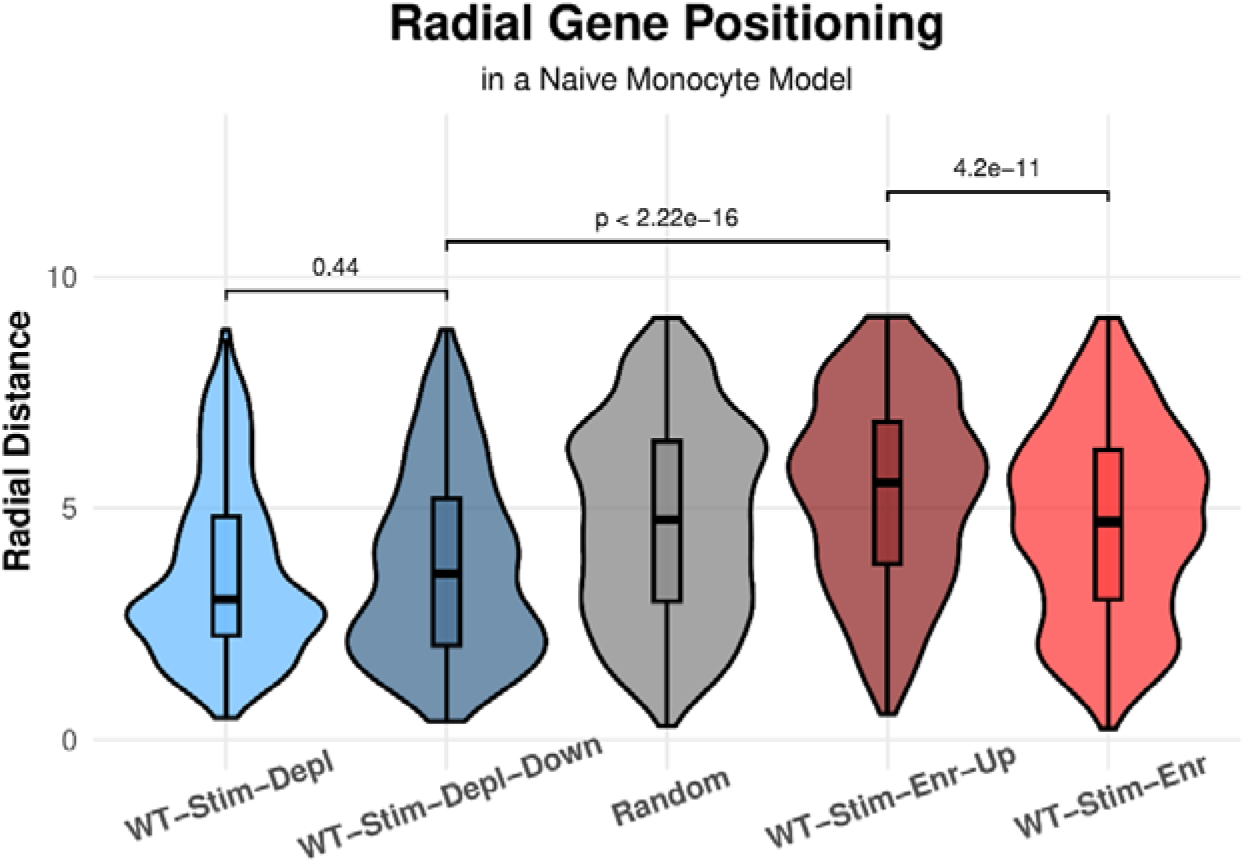
Radial distance distributions of selected gene groups on our Chrom3D^36^ model of naïve monocytes. 1000 random genes were selected from the same model. Significance of pairwise differences was assessed with Wilcoxon Rank-Sum tests.

**Supplementary Figure 3a.**
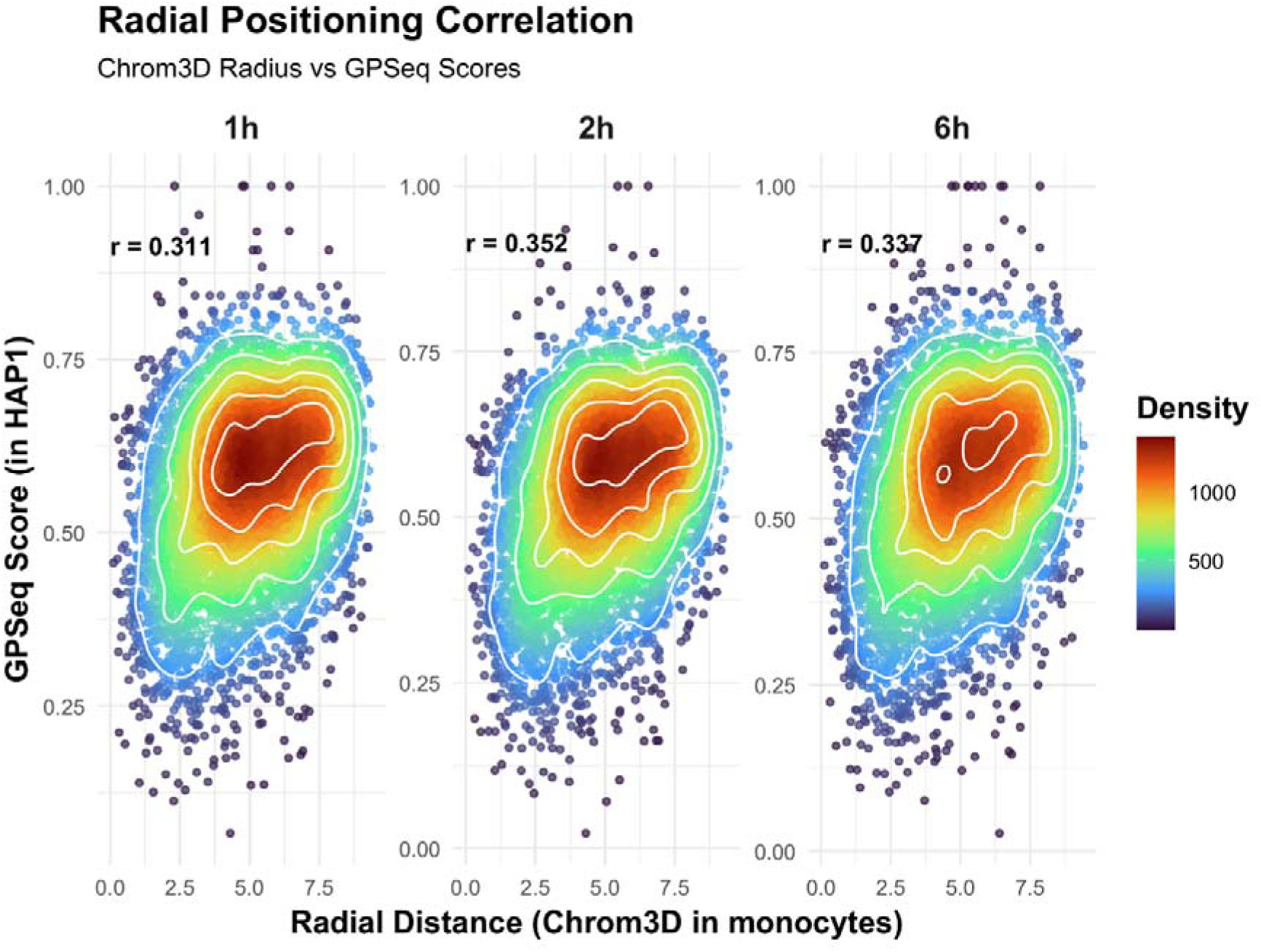
Chrom3D^36^ model calculated radial distances of monocyte TADs compared to their aggregate GPSeq Scores as assessed with GPSeq for HAP1 cells of similar myeloid origin^5^. GPSeq scores correspond to predominantly central (low scores) or predominantly peripheral regions (high scores). Increasing correlation coefficients suggest a more representative sampling of genome space at prolonged digestion times as described in^5^.

**Supplementary Figure 3b.**
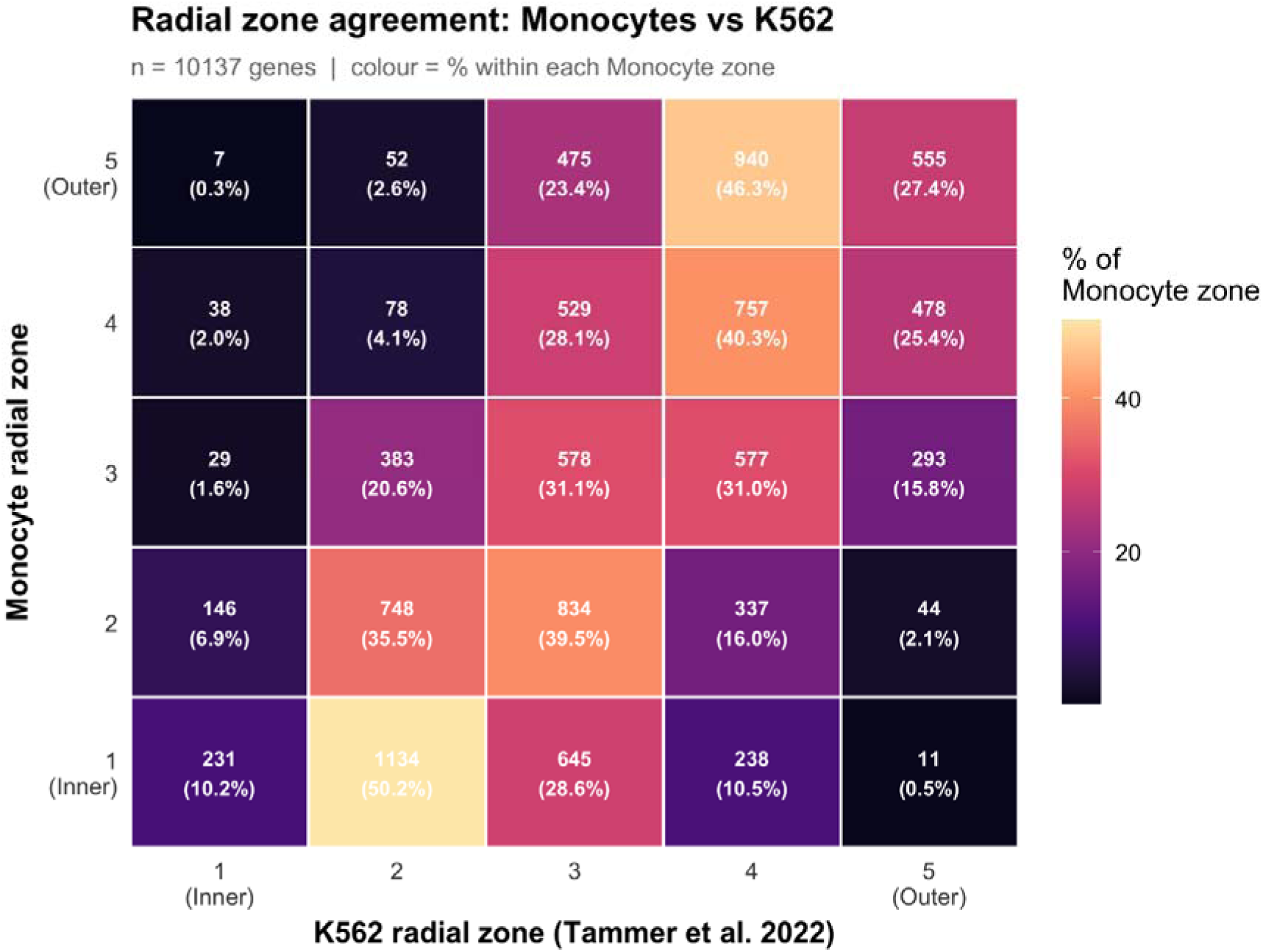
Radial zone agreements between our Chrom3D^36^ model for resting monocytes and a model produce with the same methodology for K562 cells^16^. Each cell is coloured according to the number and relative percentage of genes identified in the corresponding radial zones between the two datasets.

**Supplementary Figure 3c.**
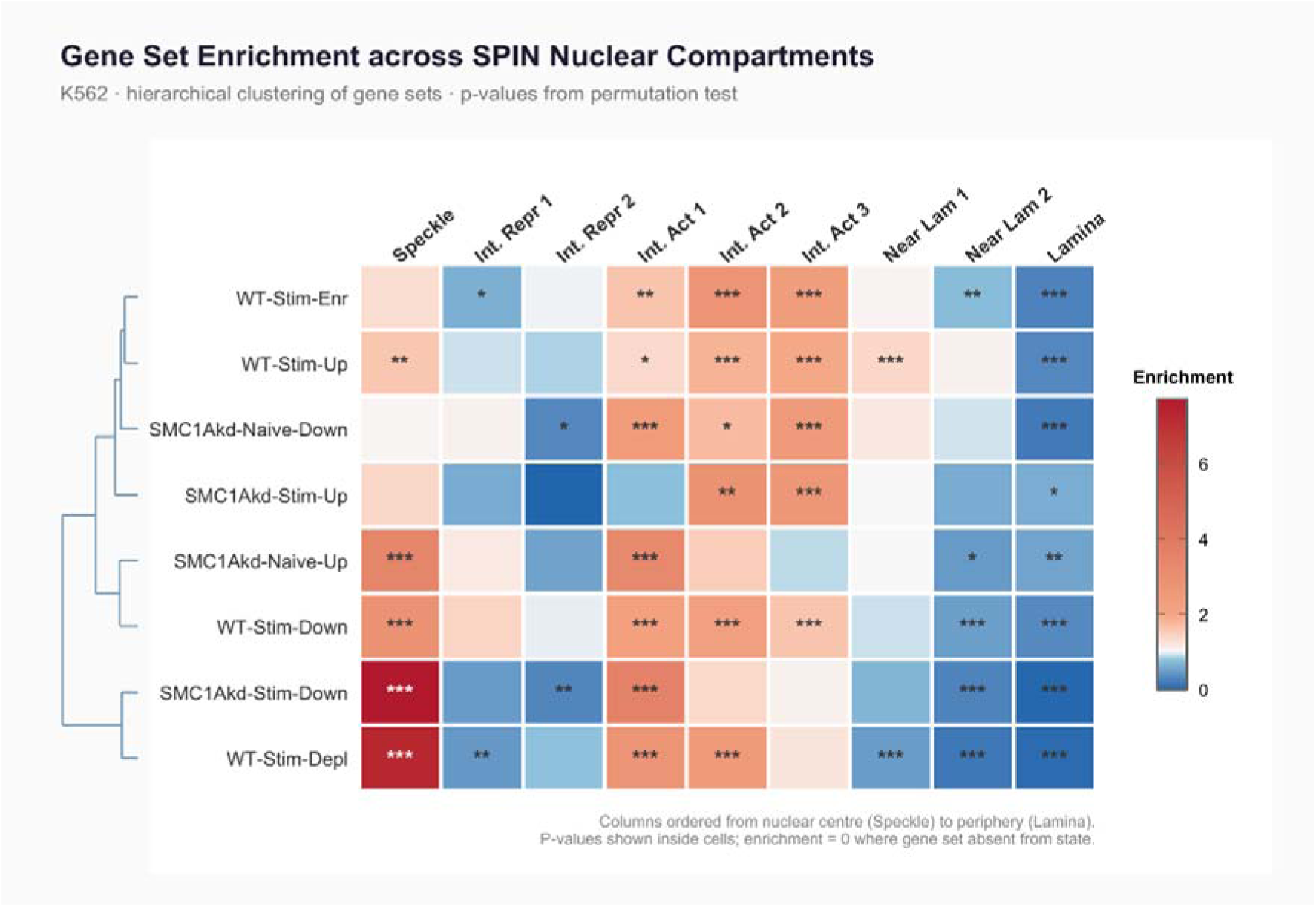
Relative enrichment of inflammatory or SMC1Akd gene groups against SPIN states in K562 cells^45^. Enrichments were calculated as observed over expected ratios of gene overlaps. P-values were assessed on the basis of a permutation test (N=1000). Stars denote significance at 0.001 (***), 0.01 (**) and 0.05 (*) levels.

**Supplementary Figure 4a.**
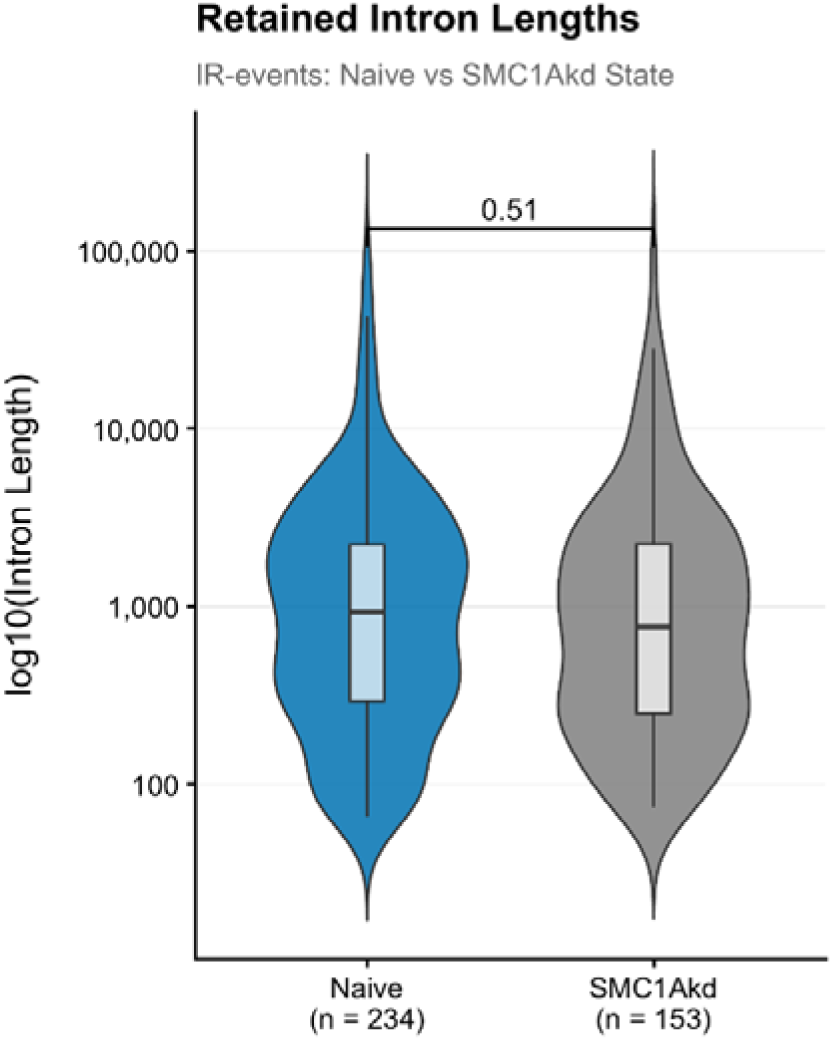
Intron sizes for introns retained in untreated or SMC1Akd monocytes. P-value of a Wilcoxon Rank Sum test.

**Supplementary Figure 4b.**
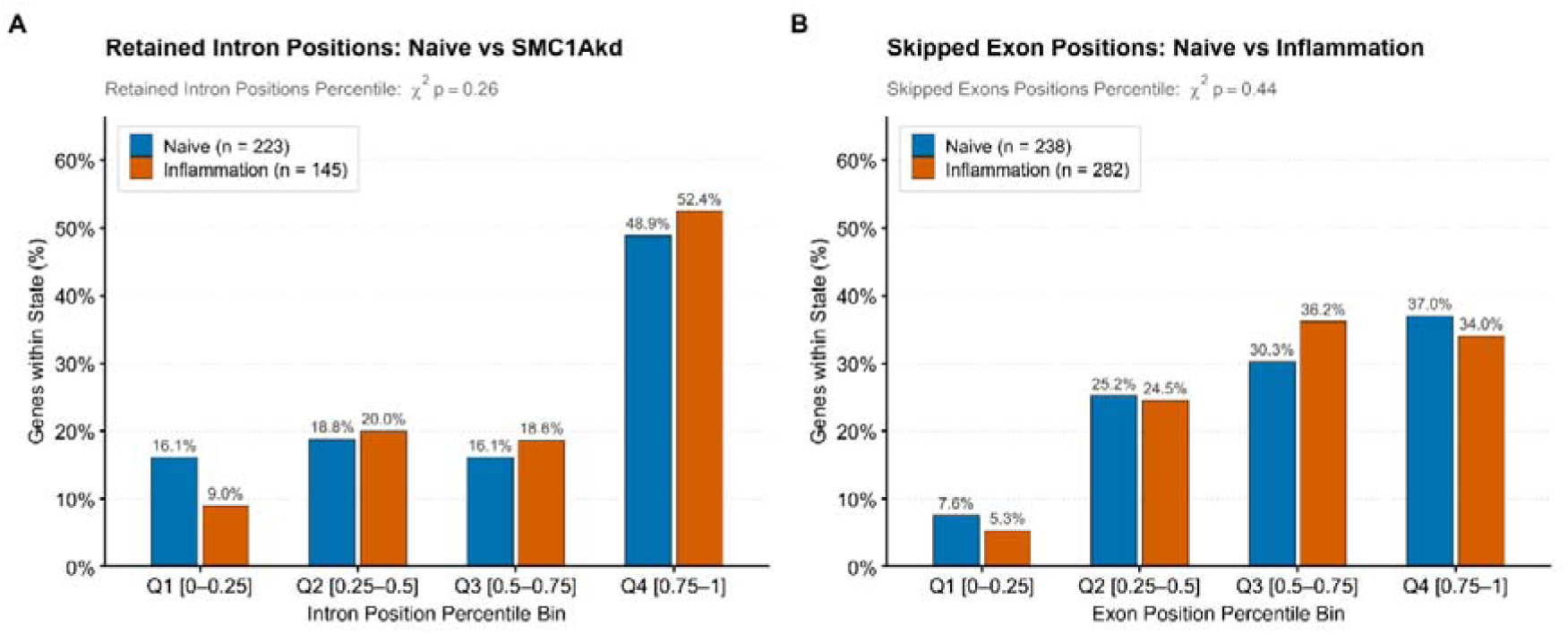
Relative position of retained introns in untreated vs SMC1Akd monocytes (left) and of skipped exons in untreated vs stimulated monocytes. Position assigned to relative quintiles corresponding to the position of the intron in the transcript, with 0-0.25 being the most 5’-proximal and 0.75-1 the most 3’-proximal bin. (Values of χ^2^-test p-values on the plots).

**Supplementary Figure 5a.**
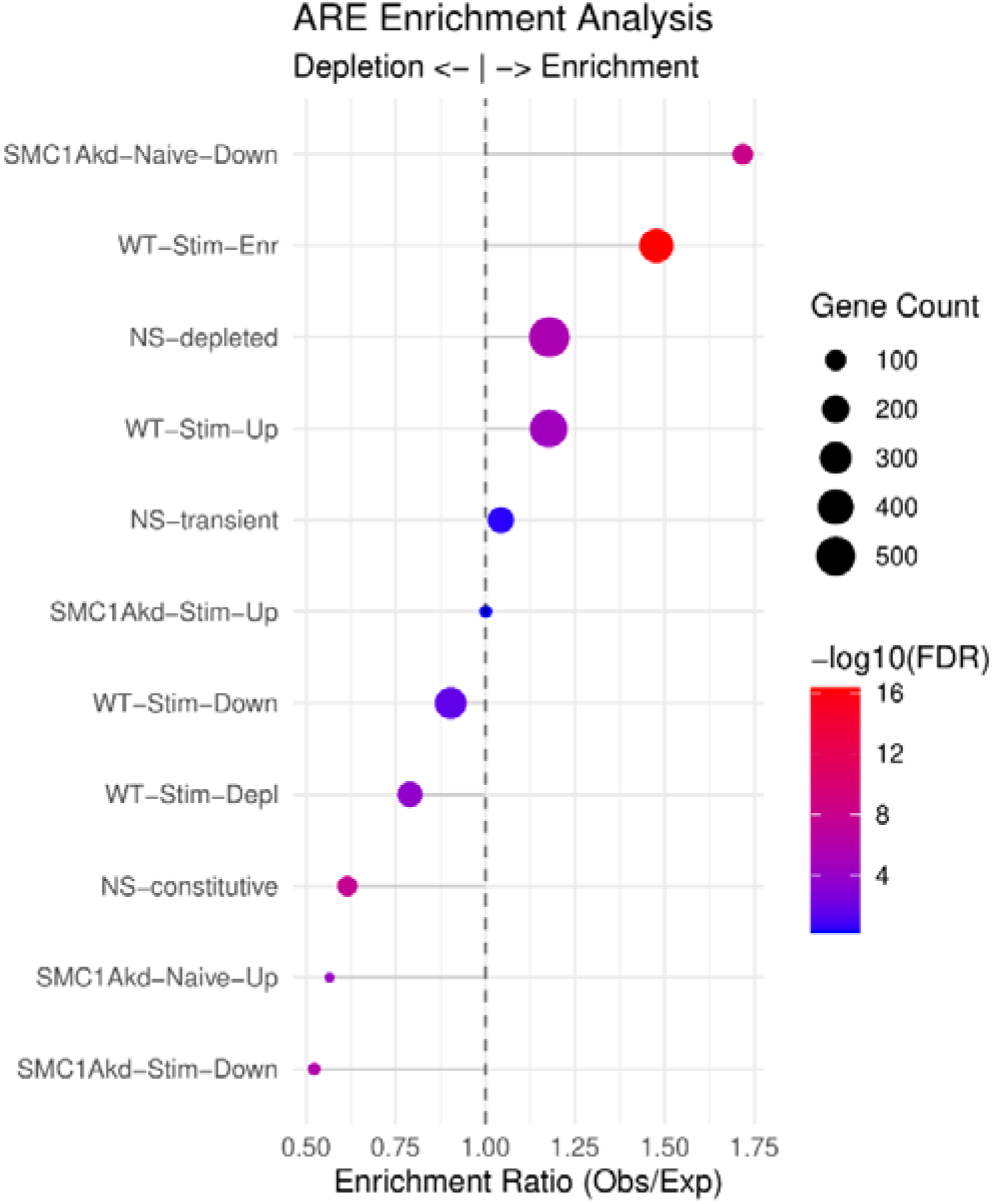
Enrichment of gene categories shown in Figure 6A in ARE elements obtained from ARED-Plus as observed/expected ratios: Values >1 correspond to enrichment, while values <1 to depletion (under-representation). Dot sizes correspond to number of genes; dot colors correspond to significance of a Fisher’s exact test (FDR corrected).

**Supplementary Figure 5b.**
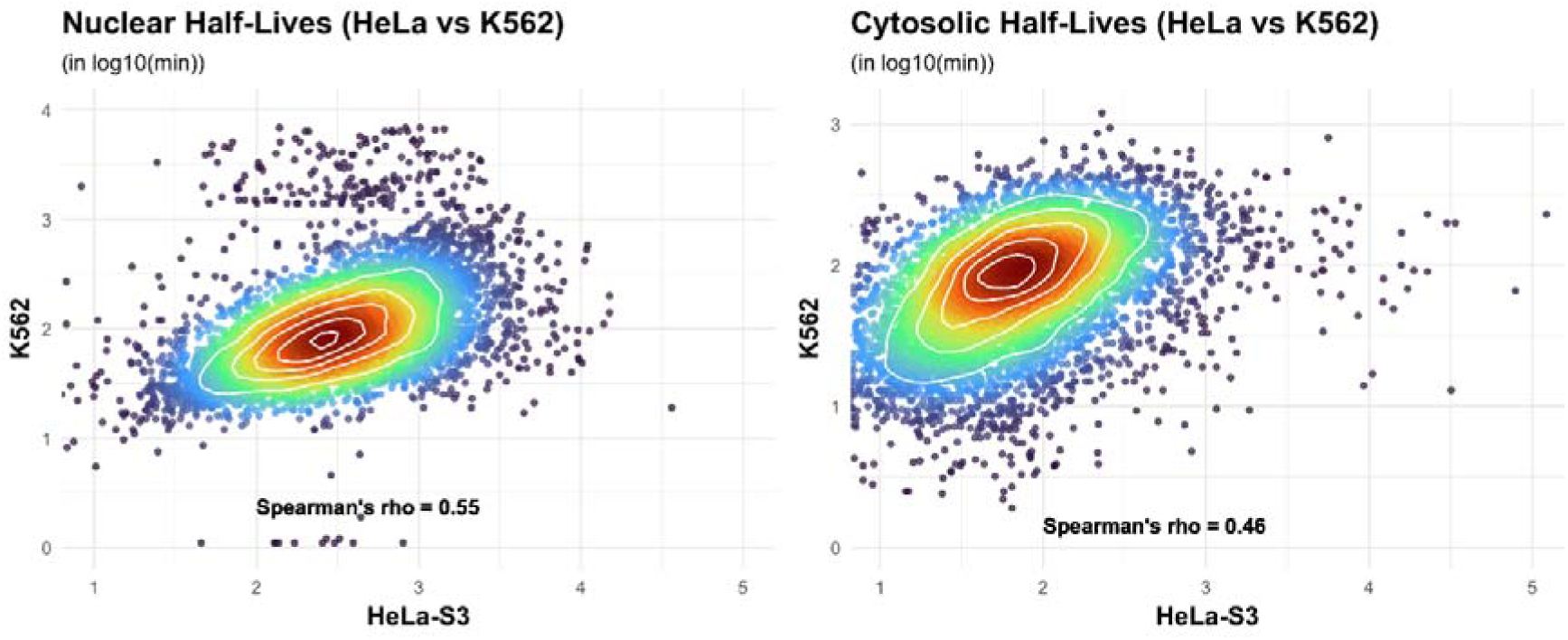
Density scatterplots and correlations of nuclear (left) and cytosolic (right) half-lives estimated with two different methods for HeLa-S3^53^ and K562 cells^56^.

**Supplementary Figure 5c.**
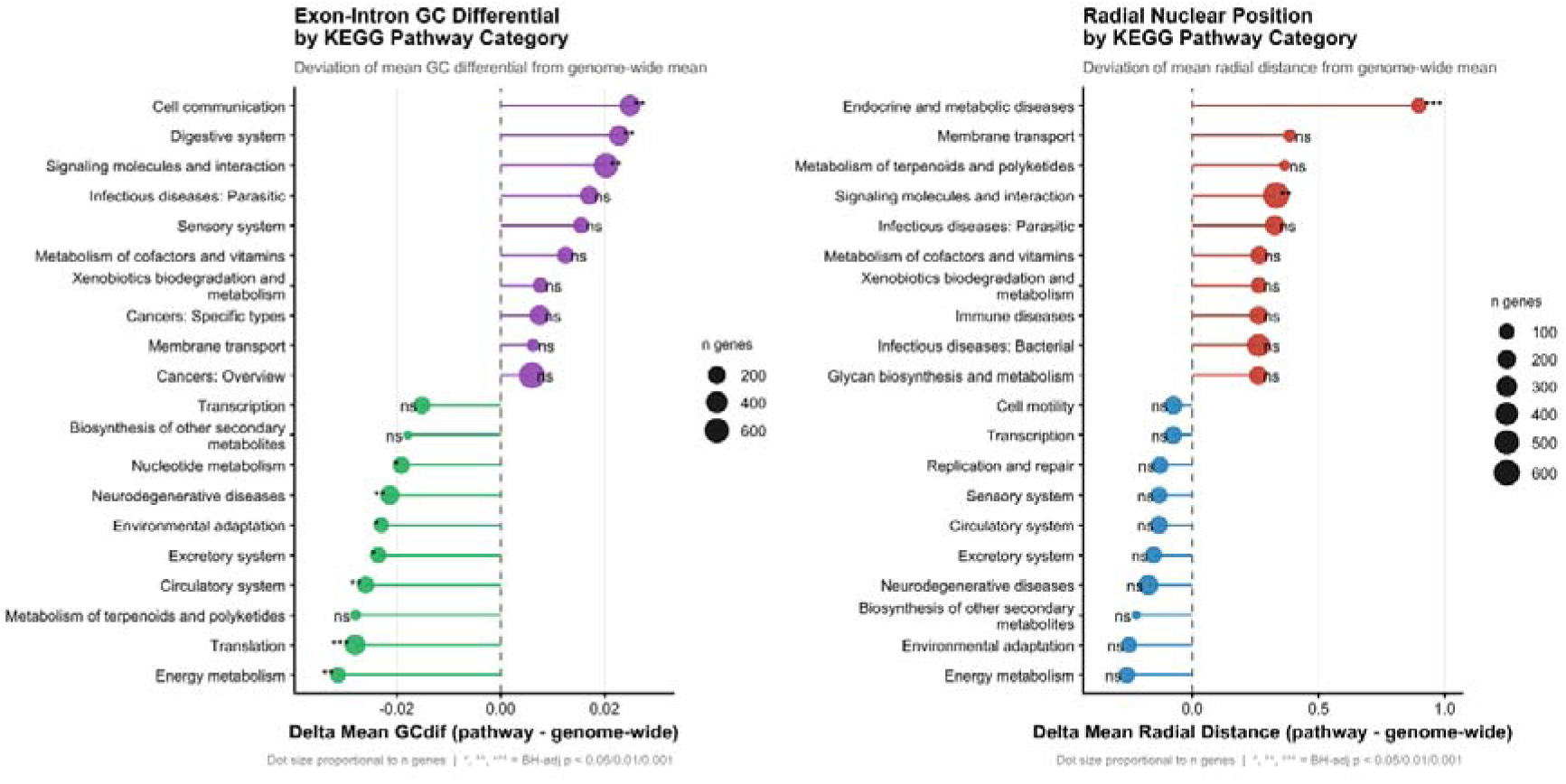
Top/Bottom 10 KEGG Pathway Categories for mean exon-intron GC% differential (left) and Radial distance estimated in our Chrom3D model of resting monocytes. Lollipop sizes correspond to gene set size. Stars denote adjusted p-values of Wilcoxon Rank Sum tests against the genome average.

